# Genesis of bursting activity: Role of inhibitory constraints on persistent Na^+^ currents

**DOI:** 10.1101/2022.02.05.479270

**Authors:** Curtis L. Neveu, Douglas A. Baxter, John H. Byrne

## Abstract

Some neurons express a plateau of spike activity greatly outlasting a stimulus, whereas others do not. This difference could be due to differential expression of ion channels (e.g., persistent Na^+^, and Ca^2+^) or similar expression of channels that differ in their properties. We compared three *Aplysia* neurons with varying capacity to generate plateau potentials: B51 with self-terminating plateaus, B64 with plateau potentials that do not self-terminate, and the regular spiking neuron B8. Our results indicate the three neurons expressed outward currents *I*_A_ and *I*_D_, voltage-gated Ca^2+^ currents *I*_CaL_ and *I*_CaR_, and persistent inward *I*_NaP_. The most notable difference observed was a larger *I*_A_, *I*_D_, and *I*_KCa_ currents in B8, and the plateau generating B64 did not express *I*_KCa_. Computational models suggest outward currents suppress and temporally constrain the plateau potential and that inward Ca^2+^ currents suppress plateau potentials when coupled with *I*_KCa_.

## INTRODUCTION

The relatively simple combination of a fast inward current and a delayed outward current can in principle produce regular neuronal spiking activity where the frequency and duration of firing is directly related to the magnitude and duration of the current stimulus (Canavier et al., 1994). However, many neurons exhibit a more nonlinear burst firing pattern driven by all-or-nothing plateau potentials, causing neurons to fire all-or-nothing bursts of activity, the duration of which can greatly outlast the stimulus (Canavier et al., 1991, 1993, 1994; Nargeot et al., 1999a,b; Plummer and Kirk, 2016). The plateau potential has increasingly become recognized as an important neurophysiological property and has been implicated in many processes such as learning and memory, hippocampal spatial learning, decision making, and rhythmic pattern generation (Bittner et al., 2017) as well as pathological conditions such as epilepsy (Stafstrom 2007; Wengert and Patel 2021). For example, plateau potentials are present in mammalian pyramidal (Bittner et al., 2017), striatal spiny projection (Carrillo-Reid et al., 2009; Plotkin et al., 2011), PreBötC respiratory (Del Negro et al., 2002), and dorsal horn neurons (Derjean et al., 2003) as well as invertebrate neurons including those from the *Aplysia* buccal ganglia (Canavier et al., 1991, 1993, 1994; Nargeot et al., 1999a,b; Plummer and Kirk, 2016). The underlying current thought to mediate many examples of plateau potentials is a persistent Na^+^ current called (*I*_NaP_) (for example see Carrillo-Reid et al., 2009) because this current has a low activation threshold, rapid activation, and a slow inactivation, but the details of this effect still remain an open question. For example, Is *I*_NaP_ present in all plateau generating neurons? Can other inward currents such as voltage-gated Ca^2+^ currents generate plateau potentials? How do outward currents such as Ca^2+^-activated K^+^ (*I*_KCa_), delayed K^+^ (*I*_D_) or A-type K^+^ (*I*_A_) channels help determine whether neurons exhibit plateau potentials?

Another question is that of complexity. Are neurons constituted parsimoniously such that regular spiking neurons contain a smaller repertoire of ion channels than plateau-generating neurons? Interestingly, *I*_NaP_ is not restricted to plateau-generating neurons. Several examples of regular firing neurons have *I*_NaP_ (Katz et al., 2018). This finding raises the question of what constrains plateau potentials and bursting activity in regular spiking neurons. Plateau-generating neurons could contain the same array of channels as regular spiking neurons, but with different properties to orchestrate plateau potentials. Although less parsimonious, containing the same array of channels with different properties could provide a neuron with greater degrees of freedom and more flexibility to adapt to different circumstances such as in learning.

The neurons in the *Aplysia* feeding system are ideal to identify the important neuronal features dictating regular spiking and plateau-generating activity patterns. Several examples of both types of neurons are present in *Aplysia*. The variety of plateau-generating neurons allows for comparisons within the same neuronal circuit. Moreover, neurons with specific firing properties can be isolated in cell culture and retain their phenotypic behavior (Lorenzetti et al., 2008; Mozziochiodi et al., 2002; Yuto et al., 2022). Here, we examined three isolated neurons in *Aplysia* that vary in their propensity to generate plateau potentials. We then characterized the major membrane currents of these neurons using two-electrode voltage clamp and pharmacologic isolation.

## RESULTS

### Neurons B51, B64, and B8 vary in their ability to generate plateau potentials

To investigate the ionic mechanisms of plateau potentials, three neurons were chosen that differ in their ability to generate plateau potentials. The B51 neuron generates a plateau potential that self-terminates after 10 – 20 s (Fig. 1A). B64 generates a plateau potential that does not self-terminate (Fig. 1A), and B8 is a regular spiking neuron unable to generate a plateau potential regardless of the amount of current injected (Fig 1A). We sought to investigate whether different ion channels mediate plateau potentials in B51 and B64 and whether these channels are more similar to each other than they are to the regular firing neuron B8? To investigate this issue, these neurons were isolated from the buccal ganglia (Fig. 1B), maintained in culture and analyzed with two-electrode voltage clamp techniques (Fig. 1C). There were noticeable differences in the morphology of cultured B51, B64 and B8 neurons (Fig. 1D).

**Figure 1.**
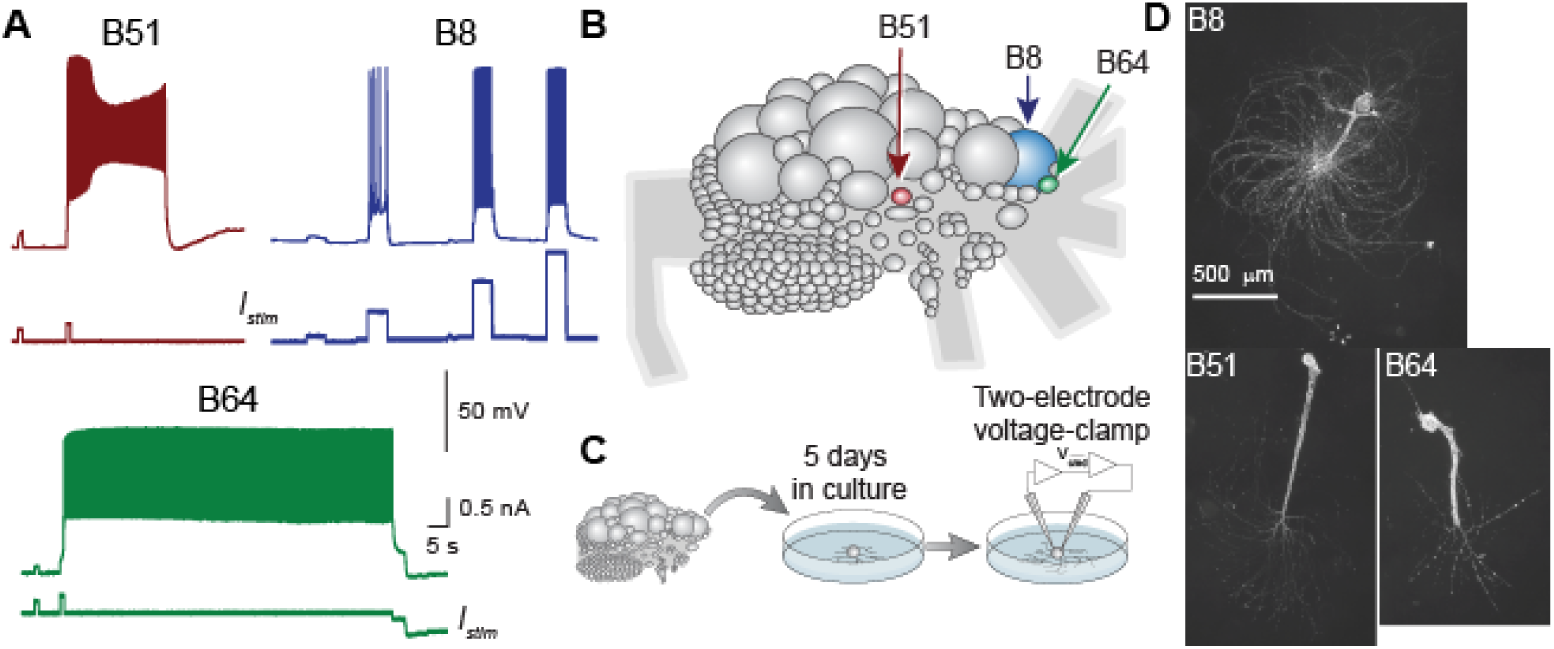
Physiological properties of bursting and pacemaker neurons. (A) Firing properties of neurons B51, B64 and B8. Activity was elicited by 1 s-duration intracellular current injection in B51 and B64, and with a 5 s-duration current injection in B8 because the large membrane time constant in this neuron. (B) Diagram of rostral surface of buccal ganglion. Neurons were identified by size and position. (C) Neurons were isolated and recorded by two electrode voltage clamp after 5 – 6 days in culture. (D) Images of cultured B51, B64, and B8 neurons.

### Outward currents of bursting and regular firing neurons

We began our analysis by characterizing the fast activating, fast inactivating *I*_A_, the relatively slower activating and inactivating *I*_D_, and the Ca^2+^ activated *I*_KCa_, which are well known to regulate action potential repolarization (for example see Carrasquillo et al., 2012; Newkirk et al., 2021; Yuan et al., 2005). ***I***_**A**_. *I*_A_ was isolated using the selective antagonist 4-aminopyridine (4-AP) (see methods). The maximum conductance g_max_ of *I*_A_ was larger in B8 compared to B51 and B64 (B51 = 1.02_(0.69, 1.1)_ µS, B64 = 0.53_(0.51,0.54)_ µS, B8 = 2.6_(1.8,3.1)_ µS) (median_(25th,75th)_) (F_(2,13)_ = 23.7, P = 0.0001; *Post hoc*, B51 vs. B64, P = 0.36; B51 vs. B8, P = 7.2×10^−4^; B64 vs. B8, P = 1.4×10^−4^) suggesting a role of *I*_A_ in suppressing plateau potentials. We then examined the slope parameter and halfway point (half-activation, half-inactivation) of the *I*_A_ steady-state voltage-gating (Eq. 12). The half-activation of *I*_A_ in B51 is shifted to more hyperpolarized potentials compared to B64 and B8 (B51 = -26.7_(−28.7, -25.0)_ mV, B64 = - 10.5_(−13.1, -6.7)_ mV, B8 = -2.5_(−11.8, -0.72)_ mV) (F_(2,13)_ = 22.5, P = 0.0001; *Post hoc*, B51 vs. B64, P = 0.0014; B51 vs. B8, P = 0.00014; B64 vs. B8, P = 0.50). The half-inactivation of *I*_A_ was also shifted to more hyperpolarized potentials in B51 compared to B64 and B8 (B51 = -55.8_(−59.4, -55.8)_ mV, B64 = -46.3_(−52.4, -44.5)_ mV, B8 = -51.0_(−51.8, -45.9)_ mV) (F_(2,13)_ = 5.94, P = 0.018; *Post hoc*, B51 vs. B64, P = 0.029; B51 vs. B8, P = 0.036; B64 vs. B8, P = 0.95). The inactivation slope parameter (Eq. 15) had a lower absolute value (corresponding to a steeper slope) in B51 compared to B8 (B51 = -4.3_(−5.0, -3.9)_, B64 = -7.4_(−8.1, -6.5)_, B8 = -9.4_(−12.2, -7.4)_) (F_(2,13)_ = 10.4, P = 0.0029; *Post hoc*, B51 vs. B64, P = 0.082; B51 vs. B8, P = 0.02 ; B64 vs. B8, P = 0.19). There was a trend in a difference in the activation slope parameter (Eq. 12) between the neurons (B51 = 5.6_(4.6, 10.7)_, B64 = 11.2_(10.6, 13.3)_, B8 = 10.1_(9.7, 11.7)_) (F_(2,13)_ = 3.18, P = 0.081). For B51, the leftward shift and steeper slope in the inactivation curve would lead to complete inactivation of *I*_A_ during the plateau. Depending on the characteristics of *I*_D_ and *I*_KCa_, the inactivation of *I*_A_ may depolarize the neuron, leading to greater activation of slower outward currents that ultimately terminate B51’s plateau. Additionally, the greater g_max_ of *I*_A_ in B8 could help prevent any plateau (see below). This greater g_max_ in *I*_A_ and the other channels can be explained by the larger size in B8 (Fig. 1 and S4). We investigated the role of morphology in plateau generation and discuss those results in a later section (see below).

***I***_**D**_. Together with *I*_A_, properties of this current could potentially explain differences in plateau potential generation between B51, B64, and B8. B8 had a larger g_max_ of *I*D compared to B51, and B64 tended to have a larger g_max_ compared to B51 (B51 = 1.6_(1.2, 2.0)_ µS, B64 = 2.9_(2.3, 3.5)_ µS, B8 = 3.5_(3.3, 4.1)_ µS) (F_(2,14)_ = 9.19, P = 0.0038; *Post hoc*, B51 vs. B64, P = 0.033; B51 vs. B8, P = 0.0034; B64 vs. B8, P = 0.43). The half-activation was not different between the three neurons (B51 = -5.0_(−6.7, -3.0)_ mV, B64 = -9.4_(−10.6, -3.1)_ mV, B8 = -5.9_(−8.9, -2.8)_ mV) (F_(2,14)_ = 0.31, P = 0.74). The slope of activation was not different between the three neurons (B51 = 9.0_(8.3, 9.1)_, B64 = 9.2_(7.3, 9.9)_, B8 = 8.6_(7.4, 9.5)_) (F_(2,14)_ = 0.07, P = 0.93). *I*_D_ did not completely inactivate for all tested command potentials. The maximum level of inactivation was compared between neurons and found to be significantly greater (lower levels of B in Eq. 15) for B8 compared to B64 (B51 = 0.36_(0.29, 0.42)_, B64 = 0.40_(0.37, 0.44)_, B8 = 0.27_(0.26, 0.31)_) (F_(2,14)_ = 5.43, P = 0.021; *Post hoc*, B51 vs. B64, P = 0.33; B51 vs. B8, P = 0.21; B64 vs. B8, P = 0.016). There was no significant difference in the half-inactivation (B51 = -10.5_(−16.9, -9.3)_ mV, B64 = -10.6_(−12.2, -2.7)_ mV, B8 = -13.4_(−15.4, -9.4)_ mV) (F_(2,14)_ = 1.57, P = 0.25) or inactivation slope (B51 = -4.5_(−4.8, -4.4)_, B64 = = -4.5_(−4.8, -4.4)_, B8 = -5.7_(−5.8, -5.4)_) (H_(2)_ = 5.42, P = 0.067). These data indicate that *I*_D_ can be activated by action potentials without being inactivated by the sustained depolarization of the plateau, and thus *I*_D_ can possibly affect the duration of the plateau. Additionally, the larger g_max_ of *I*_D_ in B8 could prevent the initiation of plateau potentials.

***I***_**KCa**_. First, we examined the contribution of *I*_KCa_ to plateau generation by blocking the influx of Ca^2+^ suppressing *I*_KCa_ activation (Fig. 3A-B). The voltage-gated Ca^2+^ channel blocker Ni^2+^ increased the duration of the plateau potential in B51 (control = 5.5_(0.0, 11.4)_ s, Ni^2+^ = 32.0_(14.4, 65.7)_ s, *paired t-test, t*_*(7)*_ = 2.882, P = 0.024). Plateaus lasting longer than 80 s were terminated artificially by negative current injection. In some cases, B51 neurons which did not express a plateau potential in control saline, exhibited a strong plateau potential in the presence of Ni^2+^. There was no significant correlation in plateau duration before and after treatment with of Ni^2+^ (*Pearson’s r* = -0.355, P = 0.38, n = 8). These data suggest that *I*_KCa_ is present in B51 and provides an inhibitory constraint on plateau potentials. We next sought to characterize *I*_KCa_ and compare its properties amongst the three neurons. *I*_KCa_ was found in B51 and B8, but not in B64 (Fig. 3D). As with the other outward currents, B8 had a markedly greater g_max_ compared to B51 (B51 = 3.5_(1.9, 4.9)_ µS, B8 = 13.6_(12.9, 27.5)_ µS, *t-test, t*_*(11)*_ = 4.52, P = 8.7×10^−4^). There was no statistically significant difference in the half-activation between B51 and B8 (B51 = 4.1_(−8.9, 8.4)_ mV, B8 = -1.7_(−4.8, 4.9)_ mV, *t-test, t*_*(12)*_ = -0.135, P = 0.90). There was also no difference in activation slope between B51 and B8 (B51 = 2.6_(0.55, 8.0)_, B8 = 7.9_(2.6, 10.7)_, *t-test, t*_*(12)*_ = 0.576, P = 0.58).

### Voltage-gated Ca^2+^channels

Two Ca^2+^ currents, *I*_CaL_ and *I*_CaR._were characterized.

***I***_**CaL**_. Similar to the outward currents, *I*_A_ and *I*_D_, the g_max_ of *I*_CaL_ was greater in B8 compared to B51 and B64 (B51 = 0.25_(0.21, 0.28)_ µS, B64 = 0.17_(0.17, 0.20)_ µS, B8 = 0.68_(0.54, 0.94)_ µS) (H_(2)_ = 10.26, P = 0.0059; *Post hoc*, B51 vs. B64, P = 0.41; B51 vs. B8, P = 0.14; B64 vs. B8, P = 0.004). The greater conductance could lead to greater influx of Ca^2+^ and greater activation of *I*_KCa_, suppressing the plateau potential in B8. There was also a leftward shift of the inactivation curve of B8 compared to B51 (B51 = -22.2_(−23.8, -19.2)_ mV, B64 = -26.5_(−27.9, -25.5)_ mV, B8 = -34.4_(−35.7, -30.0)_ mV) (H_(2)_ = 11.1, P = 0.0040; *Post hoc*, B51 vs. B64, P = 0.18; B51 vs. B8, P = 0.0026; B64 vs. B8, P = 0.27). The difference in inactivation curves would further enhances influx of Ca^2+^ in B8. There were no differences in the half-activation (B51 = -8.0_(−12.1, -5.9)_ mV, B64 = -4.7_(−8.0, 0.05)_ mV, B8 = -3.7_(−7.6, -2.5)_ mV) (F_(2,14)_ = 0.96, P = 0.41), slope of the activation curve (B51 = 7.9_(3.8, 8.8)_, B64 = 11.9_(10.1, 13.0)_, B8 = 7.9_(6.7, 13.0)_) (F_(2,14)_ = 1.71, P = 0.22), or slope of the inactivation curve (B51 = -0.99_(−4.1, - 0.91)_, B64 = -7.5_(−7.6, -6.8)_, B8 = -6.3_(−6.6, -6.0)_) (H_(2)_ = 4.5, P = 0.105).

***I***_**CaR**_. Similar to *I*_A_, *I*_D_, and *I*_CaL_, the g_max_ of *I*_CaR_ was greater in B8 compared to B51 and B64 (B51 = 0.34_(0.28, 0.38)_ µS, B64 = 0.49_(0.40, 0.63)_ µS, B8 = 2.03_(1.32, 2.5)_ µS) (F_(2,14)_ = 31.0, P = 1.12×10^-5^; *Post hoc*, B51 vs. B64, P = 0.74; B51 vs. B8, P = 1.61×10^−5^; B64 vs. B8, P = 7.61×10^−5^), which could contribute to suppression of the plateau in B8. There was also a rightward shift of the inactivation curve of B51 compared to B64 and B8 (B51 = - 18.1_(−18.4, -18.1)_ mV, B64 = -25.1_(−26.7, -22.9)_ mV, B8 = -25.3_(−26.4, -23.3)_ mV) (H_(2)_ = 9.38, P = 0.0092; *Post hoc*, B51 vs. B64, P = 0.024; B51 vs. B8, P = 0.020; B64 vs. B8, P = 0.997) and a steeper slope of the inactivation curve in B51 compared to B64 and B8 (B51 = - 1.49_(1.21, 2.18)_, B64 = -6.7_(5.54, 7.9)_, B8 = -5.95_(5.39, 7.09)_) (F_(2,14)_ = 25.5, P = 4.77×10^−5^; *Post hoc*, B51 vs. B64, P = 8.5×10^−5^; B51 vs. B8, P = 0.0002; B64 vs. B8, P = 0.82). There was no significant difference in the half-activation (B51 = 1.7_(1.4, 3.9)_ mV, B64 = 3.9_(−3.5, 4.2)_ mV, B8 = 2.0_(0.76, 3.9)_ mV) (H_(2)_ = 0.04, P = 0.98) or slope of the activation curve between the three neurons (B51 = 7.2_(2.4, 8.5)_, B64 = 7.0_(6.7, 7.3)_, B8 = 4.8_(4.4, 5.5)_) (F_(2,15)_ = 1.67, P = 0.226).

The activation curve of *I*_CaR_ was shifted to the right compared to *I*_CaL_ suggesting that *I*_CaL_ may contribute more to Ca^2+^ influx during the plateau compared to *I*_CaR_. Also, it is worth noting that the activation and inactivation curves indicate that these channels are not inactivated at resting potential become active only during neuronal activity.

### Persistent Na^+^ channel

We hypothesized that *I*_NaP_ is more pronounced in B51 and B64 compared to B8 because of *I*_NaP_ established role in maintaining sustained depolarizations. As a first step, we sought to identify a pharmacologic agent that could block *I*_NaP_ without blocking the faster *I*_Na_. Therefore, we completed a dose response curve for two compounds suggested to block *I*_NaP_, TTX and rilizule. We measured the duration of the plateau potential as an indicator for *I*_NaP_ and the amplitude of the action potential as an indicator of *I*_Na_ (Fig. 5A-B). The dose response curve indicated that 10 µM was a reasonable concentration of TTX that blocked approximately 91.2±1.3% of *I*_NaP_ while only blocking 20.6±0.7% of *I*_Na_. A similar experiment with rilizule indicated that 20 µM rilizule blocked 79.5±0.2% of *I*_NaP_ while only blocking 14.7±0.4% of *I*_Na_. However, voltage-clamp experiments with rilizule was not specific to voltage-gated sodium channels, whereas TTX was highly specific to *I*_NaP_ at 10 µM. Therefore, 10 µM TTX was selected to characterize *I*_NaP_. Surprisingly, *I*_NaP_ was not significantly different among the three neurons. There was no statistical difference in half-activation (B51 = -47.1_(−49.9, -38.1)_ mV, B64 = -39.5_(−44.3, -36.0)_ mV, B8 = -44.7_(−50.8, -42.2)_ mV) (F_(2,24)_ = 2.76, P = 0.085), slope of the activation curve (B51 = 2.2_(1.3, 3.3)_, B64 = 2.4_(1.9, 5.5)_, B8 = 2.7_(1.3, 5.5)_) (F_(2,24)_ = 0.39, P = 0.68), or g_max_ (B51 = 0.34_(0.25, 0.39)_ µS, B64 = 0.23_(0.16, 0.42)_ µS, B8 = 0.38_(0.29, 0.61)_ µS) (F_(2,24)_ = 0.87, P = 0.43). This indicates that the different propensities to generate plateau potentials amongst B51, B64 and B8 must depend on the outward currents or the voltage-gated Ca^2+^ channels.

### Computational models replicate dynamics of B51, B64, and B8

A conductance-based model of each neuron was built to investigate the ways in which these channels interact to mediate plateau potentials. For simplicity, we modeled each neuron as a single compartment. A model of each channel was constructed in agreement with the empirically measured voltage-dependency of steady-state activation and inactivation (Figs. 2-5) and estimations of the voltage-dependency of the time constants (Supplementary Fig. 2). The *I*_HCN_ current was characterized and added to B8 because of its pronounced sag potential and ability to produce rebound excitation in this neuron (Supplementary Fig. 3G). The resultant models reproduced the salient dynamics of each neuron. Specifically, the B51 model expressed a plateau potential that self-terminated, the B64 model expressed a plateau potential that did not self-terminate, and the B8 model did not express a plateau potential for any of the stimuli tested (Fig. 6D).

**Figure 2.**
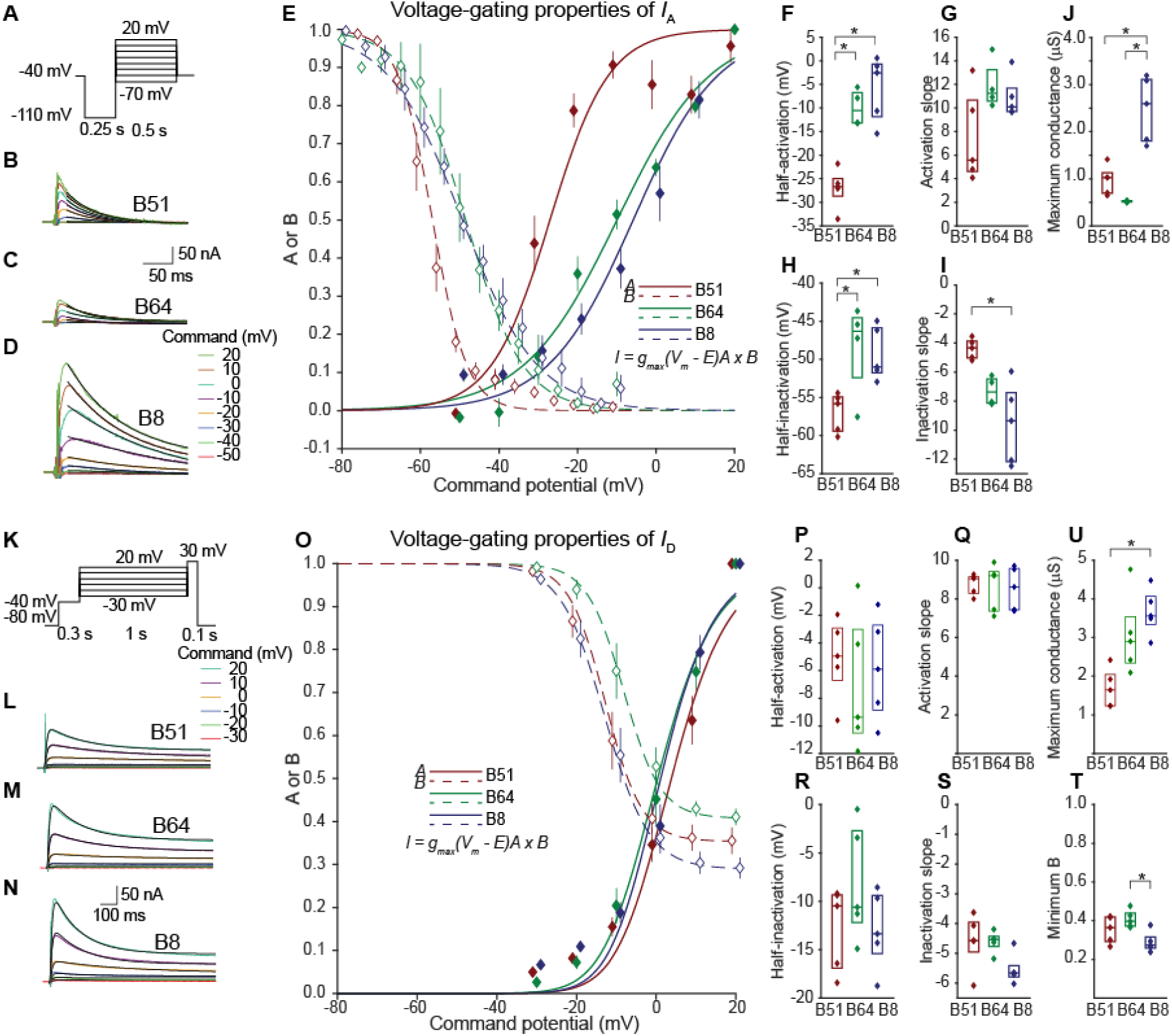
In the plateau-generating neuron B51, *I*_A_ and *I*_D_ have decreased conductances and leftward shifts in both activation and inactivation curves, as compared to the regular firing neuron B8. (A) Voltage-clamp protocol to measure *I*_A_. A 250 ms hyperpolarizing prepulse was used to de-inactivate *I*_A_. 4-AP was used to block *I*_A_ (see Methods). Inactivation of *I*_A_ was measured by a separate voltage clamp protocol (Fig. S1). (B – D) 4-AP subtracted current response indicating *I*_A_ for B51 (B), B64 (C) and B8 (D) for each command potential. A single exponential equation (Eq. 1) was fitted to a 200 ms region beginning 25 ms after start of the command pulse (black line). (E) Summary data for the inactivation (dashed line, open diamonds) and activation curves (solid line, solid diamond). The data were fit with a Boltzmann equations (Eqs. 12 and 15) with minimum A and B (minimum activation and maximum inactivation) set to 0. Data are represented as box plots. Lower and upper rectangle boundaries are the 25th and 75th percentile and the horizontal line is the median. (F – I) Summary data for the maximum conductance (Eq. 7) half-activation, activation slope, half inactivation, and inactivation slope parameters of the Boltzmann equations (Eqs. 12 and 15). Data is represented by a boxplot, median is a horizontal line and the interquartile range is represented by a rectangle. (J) Maximum conductance of *I*_A_ was estimated by extrapolating the fitted exponential curve to the beginning of the pulse and calculating conductance using Ohm’s law. (K) Voltage-clamp protocol to measure *I*_*D*_. A 300 ms prepulse at -40 mV was given to inactivate and reduce contamination by *I*_A_, making use of the lower threshold of inactivation of *I*_A_. The current responses were measured in 2 mM tetraethylammonium (TEA) to block *I*_KCa_. The final 100 ms pulse to 30 mV measured the amount of inactivation. (L – N) Current response indicating *I*_*D*_ for B51 (L), B64 (M) and B8 (N) for each command potential. The product of two exponential equations (Eq. 2) was fitted to a 1 s region beginning at the start of the command pulse (black line). (O) Summary data for the inactivation (dashed line, open diamonds) and activation curves (solid line, solid diamond) and fitted Boltzmann equations (Eqs. 12 and 15). Data represented as mean and standard error. (P – T) Summary data for the parameters of the Boltzmann equations. Data is represented by a boxplot, median is a horizontal line and the interquartile range is represented by a rectangle. (U) Maximum conductance calculated using Ohm’s law.

We next examined the window current, which is calculated using Ohm’s law and the product of the steady-state activation and inactivation curves (Eqs. 7, 12, and 15). Each current was normalized to the sum of all of the currents to display the dominant sustained current at each membrane potential (Fig. 6A-C). *I*_NaP_ was the dominant sustained current at subthreshold potentials and peaks at about -40 mV, suggesting that *I*_NaP_ mediates sustained depolarization during a plateau potential. However, a more complex set of channels mediate sustained current around -60 mV. At this potential, *I*_CaL_ seems to dominate in B51 and B64, but the outward current *I*_KCa_ was greatest at this potential in B8. Outward currents *I*_D_ and *I*_KCa_ dominated the sustained current at potentials more positive than approximately -5 mV, further supporting their role in repolarizing the membrane after an action potential.

The dynamic current responses during activation differed between the neuronal models when activated by a simulated depolarizing current injection. *I*_NaP_ was more prominent in B51 and B64, whereas there were substantially greater amounts of *I*_A_, *I*_D_, *I*_KCa_, *I*_CaL_ and *I*_CaR_ currents in B8. Any of these currents could potentially lead to the suppression of plateau potentials either directly (i.e., *I*_A_, *I*_D_, *I*_KCa_) or indirectly by *I*_CaL_ and *I*_CaR_ activation of *I*_KCa_. Therefore, we examined which currents more strongly affected the propensity to generate plateau potentials.

### Relative contributions of currents to plateau potentials

To investigate how strongly each current affects the propensity to generate plateau potentials, we varied the g_max_ (Eq. 7) of either *I*_A_, *I*_D_, *I*_KCa_, *I*_CaL_, or *I*_CaR_ while also varying the g_max_ of *I*_NaP_ (Fig. 7). Currents that were not being manipulated were kept at their control values. The g_max_ was chosen because this channel property differed the most between the three neurons. The duration of the plateau potential, measured at the end of a sufficiently strong stimulus (5 nA, 1 s simulated current injection), was used to indicate the propensity to generate a plateau potential. Increasing the g_max_ of channels that strongly suppress plateau potentials would require a commensurate increase in the g_max_ of *I*_NaP_ to maintain the propensity to generate a plateau potential. Conversely, currents that have little effect on plateau potentials would not require additional increases in *I*_NaP_. With these manipulations, we found that in B51, *I*_D_, *I*_KCa_, and *I*_CaL_ strongly suppressed the plateau potential whereas *I*_CaR_ had little effect and *I*_A_ had no effect and at all. In contrast, in B64, *I*_D_ and *I*_A_ both suppressed the plateau potential. This neuron did not have *I*_KCa_, so we artificially included the *I*_KCa_ model used in B8 and varied its g_max_ to test the effect of *I*_KCa_ on B64 plateau potentials. *I*_KCa_ strongly suppressed the plateau potential in B64. *I*_KCa_ was set to zero in B64 when testing the effects of channels other than *I*_KCa_. *I*_CaR_ did not suppress and *I*_CaL_ slightly facilitated the plateau potential in B64. The B8 model was not prone to generating plateau potentials except when g_max_ of *I*_KCa_ or *I*_D_ was much lower than control values. These results indicate that *I*_D_ and *I*_KCa_ strongly suppress plateau potentials regardless of cell type, however *I*_A_ and *I*_CaL_ only suppressed plateau potentials in particular neurons, whereas *I*_CaR_ had marginal effects on the plateau potential in all three neurons. In addition to plateau duration, we examined the voltage of the plateau potential (measured at the troughs between action potentials) and found that plateau potential heatmaps were visibly similar to plateau duration heatmaps (Supplementary Fig. 5). Interestingly, a heatmap of the action potential duration (Supplementary Fig. 6) had a more complex patterning highlighting the complex relationship between the ionic currents and shape of the action potential.

In addition to examining the voltage-dependent conductances, we also examined the passive properties in cultured B51, B64, and B8 neurons to determine their role in plateau generation. B51 had a significantly greater input resistance than B64 and B8, but B8 had a greater capacitance and membrane time constant than B51 and B64 (Supplementary Fig 4A-C). Modifying the capacitance in these model neurons only had marginal effects on plateau generation indicating that passive properties are unlikely to contribute significantly to the plateau potential (Supplementary Fig 4D).

## DISCUSSION

This study comprehensively studied three neurons that varied in their ability to produce the highly nonlinear, all-or-nothing bursting activity mediated by plateau potentials. The results indicate that regular spiking neurons like B8 can be more complex than what might be expected, containing the same conductances as in plateau generating neurons. The main difference among these neurons was greater outward currents and voltage-gated Ca^2+^ currents in B8. Interestingly, all three neurons contained similar properties of *I*_NaP_. The similar array of ion channels and increased outward current suggest that plateau potentials result not from additional channels that enable an upper depolarized state, but rather a release of inhibitory constraints like *I*_D_ and *I*_KCa_.

### Similarity of *Aplysia* channels with other systems

The steady-state activation and inactivation properties of each channel had remarkable similarity to their counterparts in mammalian systems. For example, the activation and inactivation curves of *I*_D_ agree with measurements in *X. laevis* (Tsuk et al. 2005), and *I*_CaL_ agrees with measurements of human *I*_CaL_ expressed in HEK cells (Jangsangthong et al., 2010). For *I*_CaR_, the activation curve closely matched *I*_CaR_ in cerebellar granule cells, however the *I*_CaR_ inactivation curve in *Aplysia* was shifted to the right by 20 mV compared to *I*_CaR_ in these granule cells (Randall and Tsien, 1997). In this case, *I*_CaR_ would be more effective in regulating the plateau potential in granule cells than it is in *Aplysia*.

### Voltage-gated Ca^2+^ channel’s involvement in plateau generation

The results from the current study suggest that *I*_CaL_ suppressed plateau potentials in neurons that expressed *I*_KCa_ whereas *I*_CaR_ had less of an effect on plateau potentials. In barrelette cells, blocking *I*_CaL_ also blocks plateau potentials (Lo and Erzurumlu 2002). In addition, Ni^2+^ blocked bursting in hippocampal neurons suggesting a possible role of *I*_CaR_ in plateau generation in these neurons (Metz et al., 2005). The results in barrelette and hippocampal neurons could be explained by *I*_KCa_ either not being expressed sufficiently in these neurons or being expressed in a different cellular region. Another possibility is that these voltage-gated Ca^2+^ channels enhance excitatory input and this enhanced input facilitates plateau generation. This study did not investigate the T-type Ca^2+^ channel (*I*_CaT_), which has also been implicated in bursting activity (Broicher et. al., 2007). The lower threshold of *I*_CaT_ would enable this current to play a larger role in plateau initiation, however the lower inactivation threshold would likely inactivate *I*_CaT_ for the majority of the plateau. It would be interesting to compare *I*_CaT_ role in plateau potentials in B51 or B64 in future studies.

### *I*_A_, *I*_D_ and *I*_KCa_ involvement in plateau generation

This study found that the outward currents *I*_A_, *I*_D_ and *I*_KCa_ regulated plateau potentials. The strongest regulator of plateau potentials was *I*_KCa_. In the mouse pituitary, the Ca^2+^-activated channels IK and BK had opposing effects on bursting. Inhibition of IK enhanced whereas inhibition of BK decreased bursting activity (for review see Duncan and Shipston, 2016). The delayed rectifier K^+^ has also been implicated in regulating burst firing. For example, a Kv2 blocker, guangxitoxin-1E, increased bursting in layer 5 pyramidal neurons (Newkirk et al., 2021).

### *I*_NaP_ involvement in plateau generation

This study brings additional experimental and computational support for the important role of *I*_NaP_ in mediating the plateau potential. Similar to previous studies, our results indicated that *I*_NaP_ does not fully inactivate and is activated at subthreshold voltages (Pennartz et al., 1997). In addition, computational models indicate that increasing *I*_NaP_ leads to plateau potentials.

## STAR METHODS

### RESOURCE AVAILABILITY

#### Lead contact

Further information and requests for resources and reagents should be directed to and will be fulfilled by the lead contact, John H. Byrne.

#### Materials availability

No new materials were created in this study.

#### Data and code availability

- Raw data and annotated spreadsheets are deposited on Google Drive. A link will be provided on request.
- The code for mathematical simulation is provided on ModelDB. The analysis routines were written in MATLAB are provided on GitHub at http:abc.com.
- Any additional information for reanalyzing the data is available from the Lead Contact upon request.

### EXPERIMENTAL MODEL AND SUBJECT DETAILS

#### Animals

*Aplysia californica* (100 – 120 g) were obtained from the National Resource for *Aplysia* (University of Miami, FL). Animals were housed in perforated plastic cages in aerated seawater tanks at a temperature of 15 °C and were fed ∼1 g of dried seaweed three times per week.

#### Cell culture

Culturing procedures followed those described in Brembs et al. (2002) and Lorenzetti et al. (2008). Briefly, ganglia from adult *Aplysia* were treated with Dispase® II (9.4 units/ml) (neutral protease, grade II) (Roche, Indianapolis, IL) at 35°C for ∼3 h and then desheathed. Fine-tipped glass microelectrodes were used to remove individual cells from the ganglia. Each cell was isolated and plated on poly-L-lysine (0.75 mg/ml) coated petri dishes with culture medium containing 50% hemolymph, 50% isotonic L15 (Invitrogen, Carlsbad, CA). L15 was made of 350 mM NaCl, 29 mM MgCl_2_, 25 mM MgSO_4_, 11.4 mM CaCl_2_, 10 mM KCl, 2 mM HCO_3_, streptomycin sulfate (0.10 mg/mL), penicillin-G (0.10 mg/mL), dextrose (6 mg/mL) and 15 mM HEPES. The pH of the culture medium was adjusted to 7.5. Cells were maintained for 5-6 days. Prior to recording, the culture medium was exchanged for isotonic ASW solution containing 425 mM NaCl, 12.5 mM MgSO_4_, 11.2 mM CaCl_2_, 42 mM MgCl_2_, 10 mM KCl, and 12.5 mM HEPES (pH 7.6 titrated with NaOH). In cases where voltage or Ca^2+^-gated K^+^-channels were examined N-methyl-D-glucamine (NMDG) substituted Na^+^ (425 mM). The NMDG substituted ASW (nASW) was titrated with HCl.

#### Intracellular recordings

Intracellular recordings were performed at room temperature (20-22ºC) using two fine-tipped glass microelectrodes filled with 3M potassium acetate (resistance 7-15 MΩ). One electrode was used for monitoring membrane potential and a second for injecting current. Signals were amplified using the Axoclamp-2B (Molecular Devices, Sunnyvale, CA). The holding current was passed through an AAA amplifier prior to digitization. Data were acquired at 5-10 kHz with pClamp (Version 11, Molecular Devices, Sunnyvale, CA) software and filtered with a lowpass filter using Matlab with the cutoff between 300 – 700 Hz.

Whole-cell two-electrode voltage-clamp and pharmacological techniques were used to examine the properties of voltage-gated ion channels. The voltage-clamp protocol was modified to best isolate each conductance. The pharmacological agents were dissolved in 15 mL (∼5x bath volume) and perfused at a rate of 2 mL/min. For the A-type K^+^ (*I*_A_), the membrane potential was held at -40 mV, stepped to -110 mV for 250 ms to de-inactivate the *I*_A_, then immediately stepped to potentials ranging from -50 mV to 20 mV for 500 ms. The interval between each voltage-clamp stimulus (interstimulus interval, ISI) was set to 4 s. This protocol was carried out in two conditions: 200 µM 4-aminopyridine (4-AP) a concentration that does not block *I*_A_ and in 10 mM 4-AP, a concentration that inhibits *I*_A_. The low concentration of 200 µM 4-AP was chosen to reduce contamination of *I*_KCa_. Both cases used nASW containing 10 mM Ni^2+^ and 100 µM Cd^2+^ to block voltage-gated Ca^2+^ channels and *I*_KCa_. The current responses during the two pharmacological conditions were subtracted to isolate *I*_A_. For the delayed K^+^ channel (*I*_D_), the membrane potential was held at -80 mV, stepped to -40 mV for 300 ms to inactivate *I*_A_, then immediately stepped to potentials ranging from -30 mV to 20 mV for 1 s. The ISI was set to 60 s. *I*_D_ was measured in nASW containing 2 mM tetraethylammonium (TEA) and 10 mM Ni^2+^ and 100 µM Cd^2+^ to block *I*_KCa_ and voltage-gated Ca^2+^ channels. For *I*_KCa_, the membrane potential was held at -80 mV, stepped to - 30 mV for 300 ms to inactivate *I*_A_, then immediately stepped to potentials ranging from - 30 mV to 20 mV for 500 ms. The ISI was set to 3 s. This protocol was carried out in control nASW and nASW containing 2 mM TEA, a concentration that selectively inhibits *I*_KCa_ (Baxter and Byrne, 1989). The current responses during the two pharmacological conditions were subtracted to isolated *I*_KCa_. For the L-type Ca^2+^ channel (*I*_CaL_), the membrane potential was held at -80 mV and stepped to potentials ranging from -40 mV to 20 mV for 250 ms. The ISI was 30 s. This protocol was repeated in control nASW and nASW containing the *I*_CaL_ inhibitor nifedipine (10 µM). Both conditions contained 10 mM 4-AP and 50 mM TEA to block *I*_A_, *I*_D_, and *I*_KCa_. The current responses during the two pharmacological conditions were subtracted to isolate *I*_CaL_. For the R-type Ca^2+^ channel (*I*_CaR_), the membrane potential was held at -80 mV and stepped to potentials ranging from -40 mV to 20 mV for 250 ms. The ISI was set to 30 s. This protocol was repeated with a 300 ms prepulse to -10 mV to inactivate the *I*_CaR_. Both protocols were done in the presence of nASW containing 10 µM nifedipine, 10 mM 4-AP and 50 mM TEA to block *I*_CaL_, *I*_A_, *I*_D_, and *I*_KCa_. The current responses elicited by the protocol without the prepulse were subtracted from the current responses with the prepulse to isolated *I*_CaR_. For the persistent Na^+^ (*I*_NaP_), the membrane potential was held at -100 mV, stepped to -40 mV for 20 ms to inactivate the *I*_Na,fast_ channel, then immediately stepped to potentials ranging from -60 mV to -20 mV for 1 s. The ISI was set to 30 s. This protocol was repeated in ASW containing the voltage-gated Na^+^ inhibitor tetrodotoxin (10 µM TTX). The current responses during the two pharmacological conditions were subtracted to isolate *I*_NaP_.

#### Fitting equations to current responses

All automated curve fitting was done by the built-in MATLAB function lsqcurvefit using the default ‘trust-region-reflective’ or the Levenberg-Marquardt least squares algorithms. *I*_A_ current traces were fit with the following single-exponential equation:

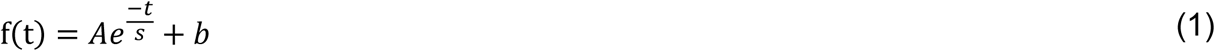

where *A* is a positive value representing the amplitude of the current, *s* is the time constant of inactivation and *b* is a baseline offset. This curve was fitted to a 200 ms region starting 25 ms after the start of the pulse for each command potential. The curve was extrapolated backwards 25 ms to obtain the initial current at the start of the pulse.

*I*_D_ current traces were fit with the following double-exponential equation:

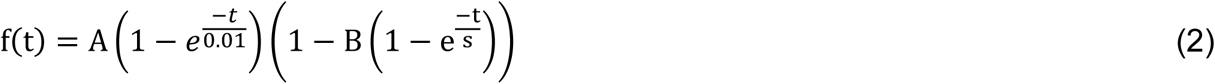

where the parameter *A* is the maximum current and is a value limited to between 1 and 1.5 times the peak of the current trace. The peak for *I*_D_ was estimated by taking the 99^th^ percentile of the current trace. To reduce the number of fitting parameters, the activation time constant was fixed at 0.01 s, a value which provided the best fit for the data. The parameter *s* is the time constant of inactivation limited to a value greater than 0.16 s. *B* is the amount of inactivation and was determined by a voltage step to 30 mV following a 1 s step to various potentials (Fig. 2K and Supplemental Fig. 1B). The second term of Eq. 2-4 is equal to B_∞_ in Eq. 15 when time is carried out to infinity. The current response measured at 18.7 ms after the start of the 30 mV pulse was normalized to the current measured with the prepulse that resulted in no inactivation (−30 mV). As for *I*_A_, *I*_CaL_, and *I*_CaR_ the inactivation data were fit with Eq. 5. The values returned from the fitted equation are inserted for *B* in Eq. 2.

*I*_KCa_ current traces were fit with the following triple exponential equation:

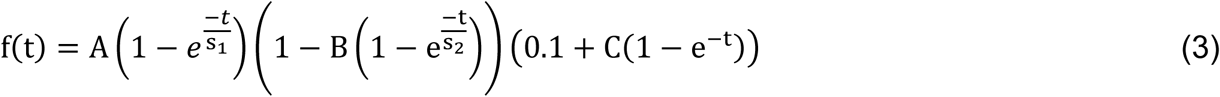

where *A* is limited to a positive value. *B* is limited to between 0 and 1. For Eq. 3, *B* was not directly measured using a separate voltage-clamp protocol. The parameters *s*_1_ and *s*_2_ are the activation and inactivation time constants respectively. The third term of Eq. 3 describes the regulation of the *I*_KCa_ by Ca^2+^. The value 0.1 is the initial value. The parameter *C* is the value of Ca^2+^ saturation.

*I*_CaL_ and *I*_CaR_ current traces were fit with the following double-exponential equation:

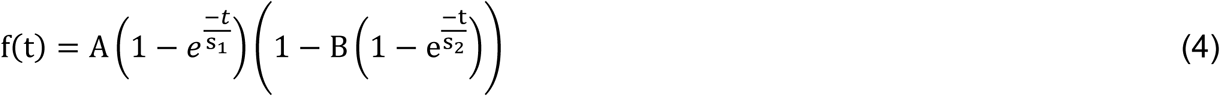

where *A* is the maximum current and limited to a value between -100 and 0 nA, which exceeded the largest current measured for these channels. *s*_1_ is the activation time constant and limited to between 0.002–0.05 s for *I*_CaL_ and 0.001–0.05 s for *I*_CaR_, *s*_2_ is the inactivation time constant and limited to between 0.02–0.5 s for *I*_CaL_ and 0.025–0.5 s for *I*_CaR_. *B* is the amount of inactivation and was measured by a separate voltage clamp protocol. A 1 s prepulse to command potentials from -70 to -10 mV was followed by a 200 ms pulse to 0 mV to measure the amount of inactivation. A single exponential of the same form as Eq. 1 was used to fit the current response to the 30 mV voltage command and extrapolated back to the start of the pulse. *B* was equal to 1 minus the extrapolated current divided by the current at -30 mV.

*I*_NaP_ current traces were fit with the following double-exponential equation:

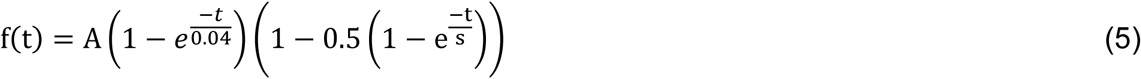

where *A* is the maximum activation and limited to between 0 and -30 nA. The value 0.04 is the activation time constant. Initial fittings indicated that the amount of inactivation was approximately 0.5 for all three neurons in the range of potentials used to measure *I*_NaP_ so we fixed the value to 0.5 for simplification and reduce the degrees of freedom of the fitting algorithm. *s* is the inactivation time constant limited to values between 0.2 and 5 s.

#### Computational model

Conductance-based models for B51, B64, and B8 were generated and simulations conducted with the Simulator for Neural Networks and Action Potentials (SNNAP, version 8.1; Baxter and Byrne 2007). The three neurons were modeled as isolated compartments with no synaptic (electrical or chemical) connections between them. The membrane potential was determined by the following differential equation:

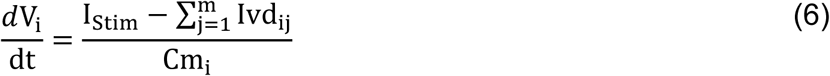

where Cm_i_ is the membrane capacitance, I_Stim_ is the extrinsic stimulus current, Ivd_ij_ denotes voltage dependent current. Each voltage dependent current was determined by the following equation:

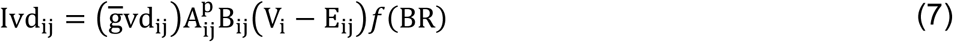

where gvd_ij_ is the maximum conductance, A_ij_ represents the voltage and time dependent activation, B the voltage and time dependent inactivation, V_i_ the membrane potential, and E_ij_ the equilibrium potential. *f*(BR) denotes regulation by a Ca^2+^ ion pool. *I*_KCa_ is regulated by two Ca^2+^ pools. The first Ca^2+^ pool that regulates *I*_KCa_ represents a constant occupied Ca^2+^ binding site of *I*_KCa_ and its regulation is constant at 0.4. The second Ca^2+^ pool is filled by voltage-gated Ca^2+^ channels. This pool’s regulation of *I*_KCa_ is determined by the following equations:

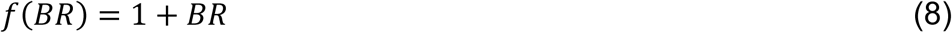

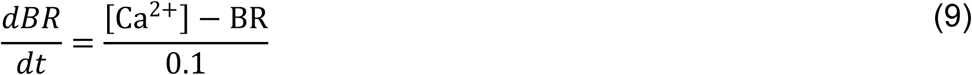

Where 0.1 is the time constant. The concentration of Ca^2+^ is determined by the following equation.

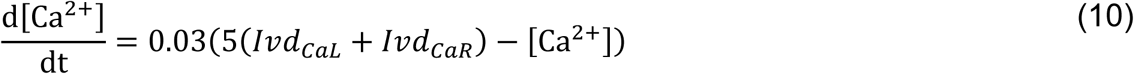

The activation A_ij_ is determined by the following Boltzmann-type equations:

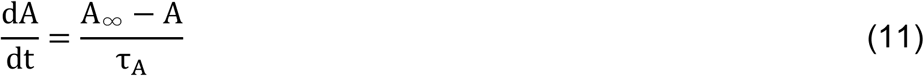

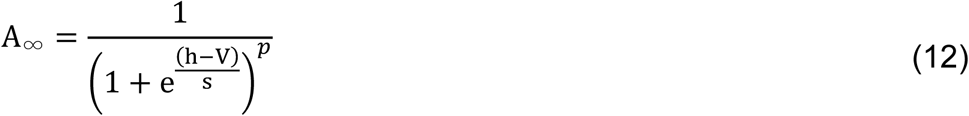

where A is the level of activation, h is the voltage of the inflection point (half-activation), V is the voltage, and s is the slope parameter. We used a Matlab curve fitting algorithm to estimate the parameters for this equation. The maximum current (*A* in Eqs. 1-5) was converted to conductance using Ohm’s law. The reversal potentials were -70 mV for *I*_A_ and *I*_D_ , -80 mV for *I*_KCa_, and 60 mV for *I*_CaL_, *I*_CaR_, *I*_Na_ and *I*_NaP_. To obtain the activation level, the conductance values at each potential were normalized to the maximum conductance of all membrane potentials measured for that cell. The activation data for each cell was fit with Eq. 12 (solid diamonds, solid lines in Figs. 2E, 2O, 3G, 4E, 4O, and 5G). For the voltage-gated Ca^2+^ channels, *h* was limited to between -50 and 10 mV and *s* was limited to between 0 and 20.

**Figure 3.**
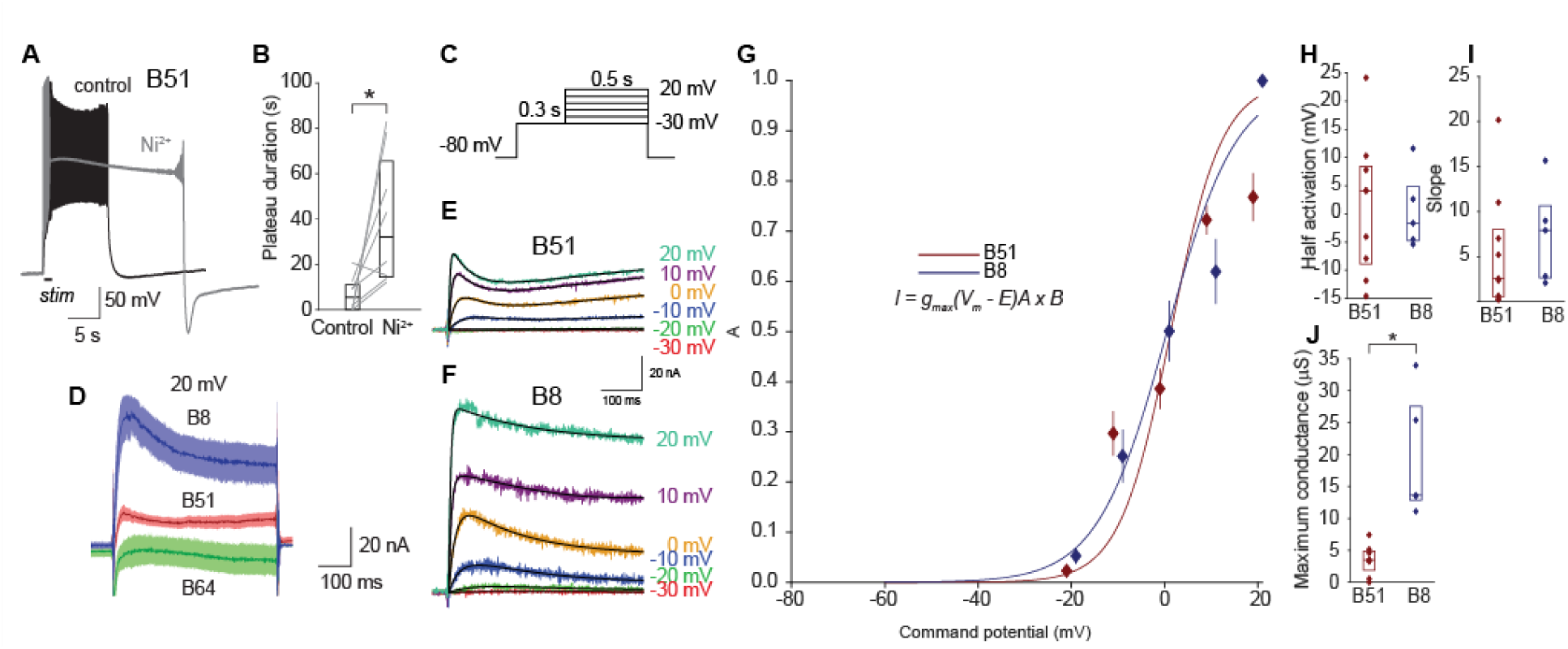
*I*_KCa_ properties. (A) Example of application of Ni2+ increasing the duration of the plateau potential of B51. (B) Summary data of effects of Ni2+ on lengthening the plateau potential of B51. Duration was measured as the amount of time above -70 mV after the end of the stimulus. Data is represented by a boxplot, median is a horizontal line and the interquartile range is represented by a rectangle. (C) Voltage-clamp protocol to measure *I*_KCa_. The current responses in control were subtracted from responses in 2 mM TEA to isolate *I*_KCa_. (D) Summary data for current responses during step to 20 mV. *I*_KCa_ can be seen for B8 and B51 but not B64. (E – F) TEA subtracted current response indicating *I*_KCa_ for B51 (E) and B8 (F) for each command potential. *I*_KCa_ was not found in B64. The product of three exponentials (activation, inactivation, and slow Ca2+ buildup, Eq. 3) was fitted for a 500 ms region beginning 1 ms after the start of the command pulse (black line). (G) Summary data for the activation curves (solid line, solid diamond) and fitted Boltzmann equations (lines, Eq. 12). Data are represented as mean and standard error. (H – I) Summary data for the parameters of the Boltzmann equations. Data is represented by a boxplot, median is a horizontal line and the interquartile range is represented by a rectangle. (J) Maximum conductance calculated using Ohm’s law.

**Figure 4.**
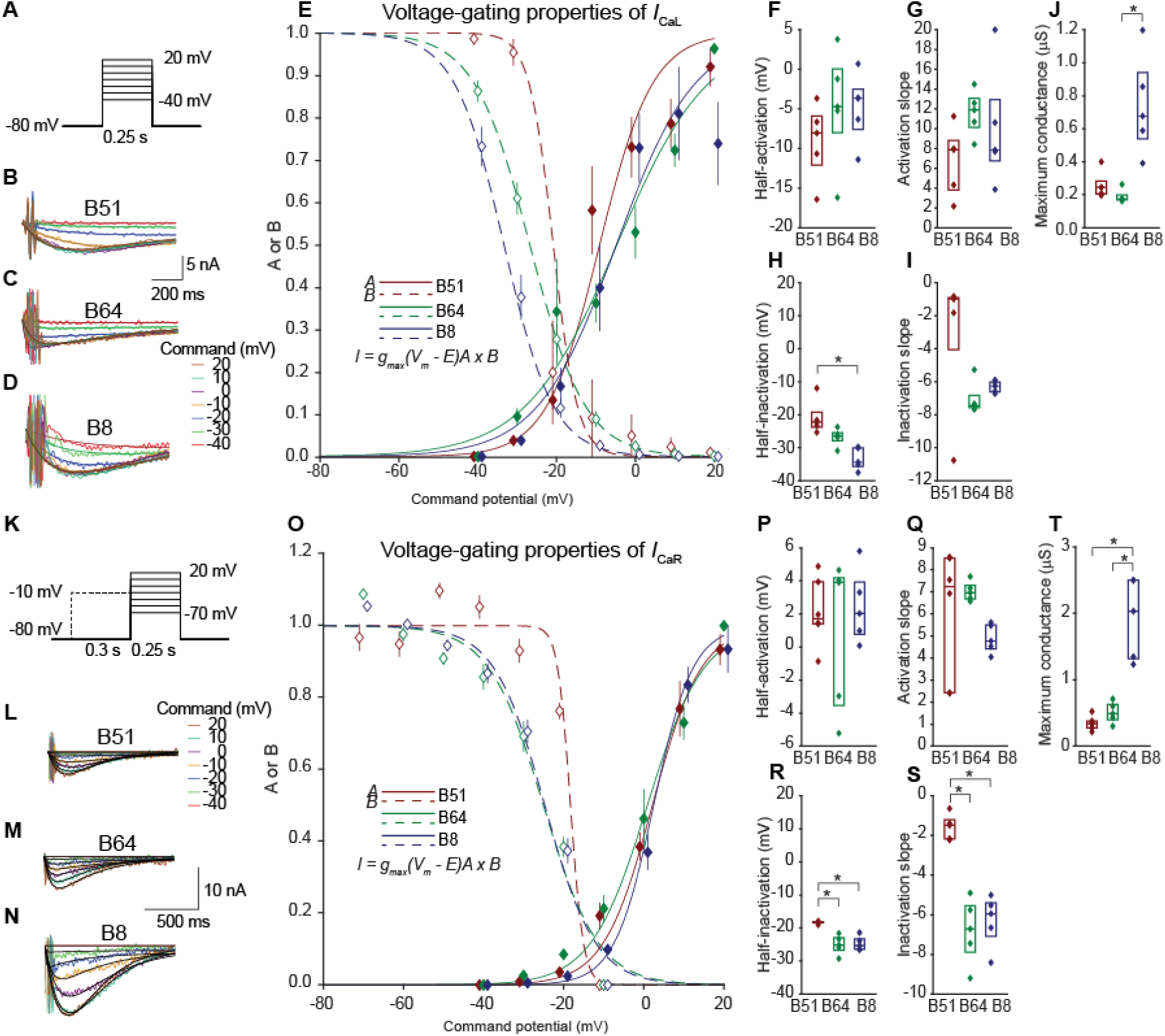
Steady-state properties of *I*_CaL_ and *I*_CaR_ indicate greater conductances in the pacemaker neuron B8 and rightward shifts in the activation and inactivation curves of B51. (A) Voltage-clamp protocol to measure *I*_CaL_. The current responses in control were subtracted from responses in 10 µM nifedipine to isolate *I*_CaL_. Inactivation for *I*_CaL_ was measured by a separate voltage clamp protocol (Fig. S2). (B – D) Subtracted current responses indicating *I*_CaL_ for B51 (B), B64 (C) and B8 (D) for each command potential. A double exponential equation (Eq. 4) was fitted to a 100 ms region beginning 2 ms after start of the command pulse (black line). (E) Summary data for the inactivation (dashed line, open diamonds) and activation curves (solid line, solid diamond). The data were fit with a Boltzmann equation (Eqs. 12 and 15) with minimum A and B (minimum activation and maximum inactivation) set to 0. Data are represented as mean and standard error. (F – I) Summary data for the parameters of the Boltzmann equations. Data is represented by a boxplot, median is a horizontal line and the interquartile range is represented by a rectangle. (J) Maximum conductance of *I*_CaL_ was calculated using Ohm’s law. (K) Voltage-clamp protocol to measure *I*_CaR_. A 300 ms prepulse at -10 mV was given to inactivate *I*_CaR_. Protocols were administered in the presence of nifedipine to block CaL. The current responses without prepulse were subtracted from responses with the prepulse to isolate *I*_CaR_. Inactivation for *I*_CaR_ was measured by a separate voltage clamp protocol (Fig. S2). (L – N) Subtracted current response indicating *I*_CaR_ for B51 (A), B64 (B) and B8 (C) for each command potential. A double exponential equation (Eq. 4) was fitted to a 120 ms region beginning 8 ms after the start of the command pulse (black line). (O) Summary data for the inactivation (dashed line, open diamonds) and activation curves (solid line, solid diamond) and fitted Boltzmann equations (lines, Eqs. 12 and 15). Data represented as mean and standard error. (P – T) Summary data for the parameters of the Boltzmann equations. Data is represented by a boxplot, median is a horizontal line and the interquartile range is represented by a rectangle. (U) Maximum conductance calculated using Ohm’s law.

**Figure 5.**
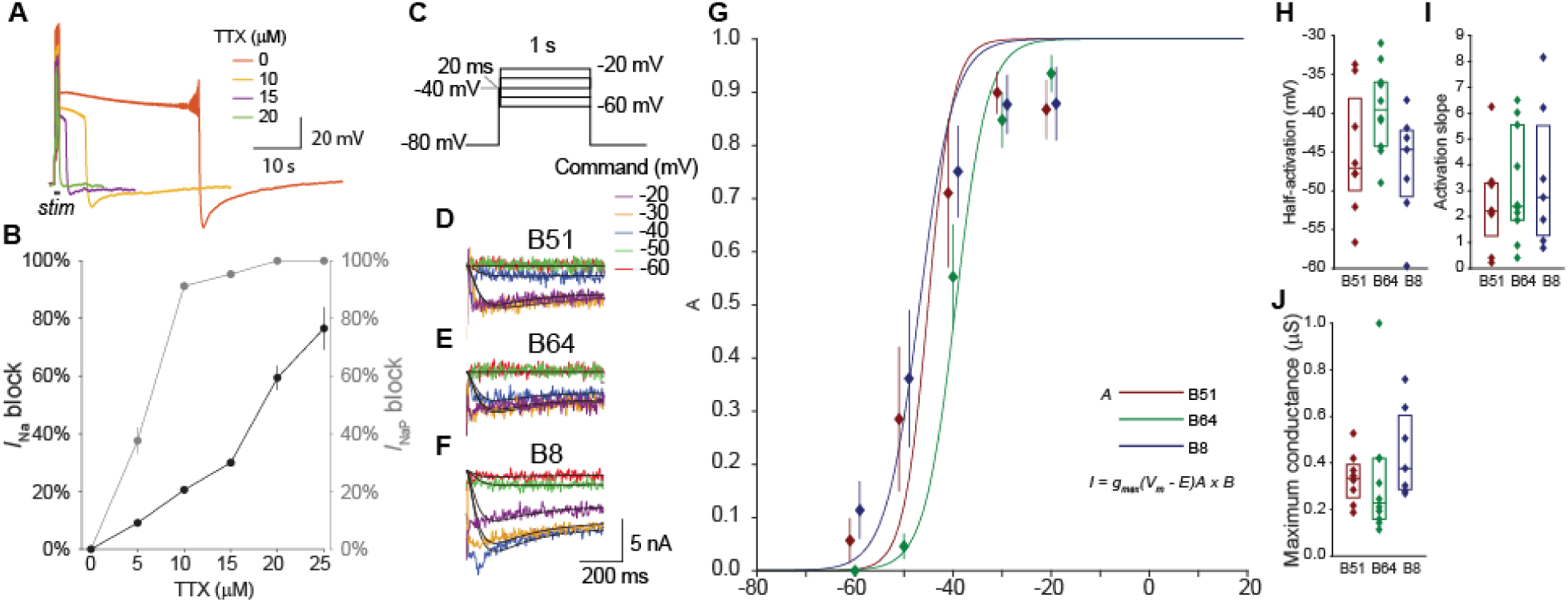
*I*_NaP_ expressed in both bursting and pacemaker neurons has similar properties. (A) Example neuron showing the reduction in plateau duration by TTX. (B) Dose response curve of TTX. The block of *I*_Na_ was measured as the decrease in action potential amplitude, whereas the block of *I*_Nap_ was measured by the percent decrease in the amount of time above -70 mV after the end of the stimulus. (C) Voltage-clamp protocol to measure *I*_Nap_. A 20 ms step to -40 mV was used to reduce contamination by *I*_Na_. The current responses in control were was subtracted from responses in 10 µM TTX to isolate *I*_Nap_. (D – F) TTX subtracted current response indicating *I*_Nap_ for B51 (D), B64 (E) and B8 (F) for each command potential. A double exponential equation (Eq. 5) was fitted for a 640 ms region beginning 9 ms after the start of the command pulse (black line). (G) Summary data for the activation curves (solid line, solid diamond) and fitted Boltzmann equations (lines, Eq. 12). Data represented as mean and standard error. (H – I) Summary data for the parameters of the Boltzmann equations. (J) Maximum conductance calculated using Ohm’s law.

**Figure 6.**
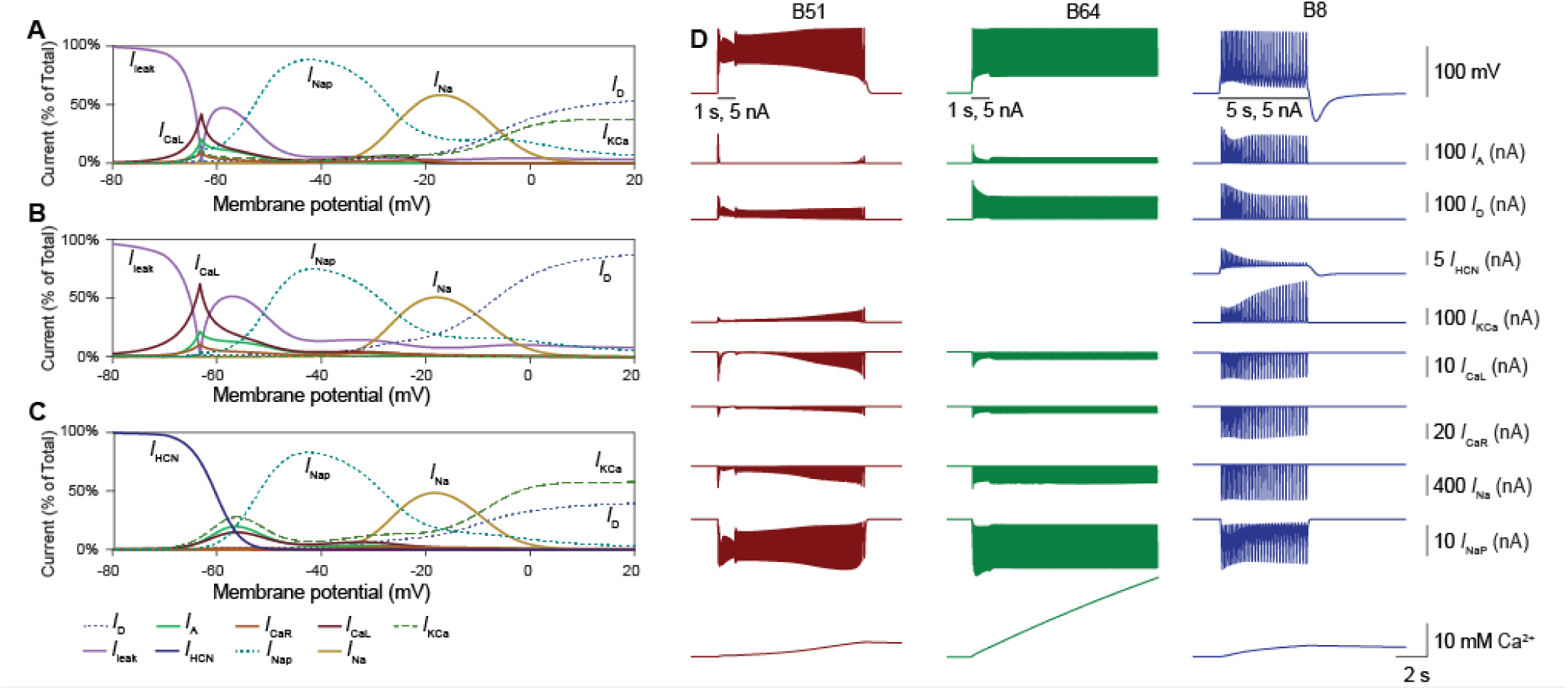
Simulations indicated outward currents *I*_KCa_, *I*_A_, and *I*_D_ terminated plateau potentials and could suppress the initiation of plateau potentials. (A – C) Plots showing the dominant steady-state current for each membrane potential. *I*_leak_ and *I*_HCN_ dominate at holding potential (−80 mV), *I*_NaP_ dominates at subthreshold depolarization, *I*_Na_ dominates at depolarized potentials, *I*_D_ and *I*_KCa_ dominate at depolarized potentials. Currents such as *I*_A_ completely inactivate at all potentials so are not observed. (B) Current responses and Ca2+ pool during simulated activity triggered by a 5 nA current injection (1 s for B51 and B64, 5 s for B8). B8 has a pronounced hyperpolarization sag potential. We confirmed the presence of *I*_HCN_ by Cs^+^ and ZD7288 and then characterized *I*_HCN_ in B8 (Supplementary Fig. [not yet added]). *I*_HCN_ was then substituted it for *I*_Leak_ in B8.

**Figure 7.**
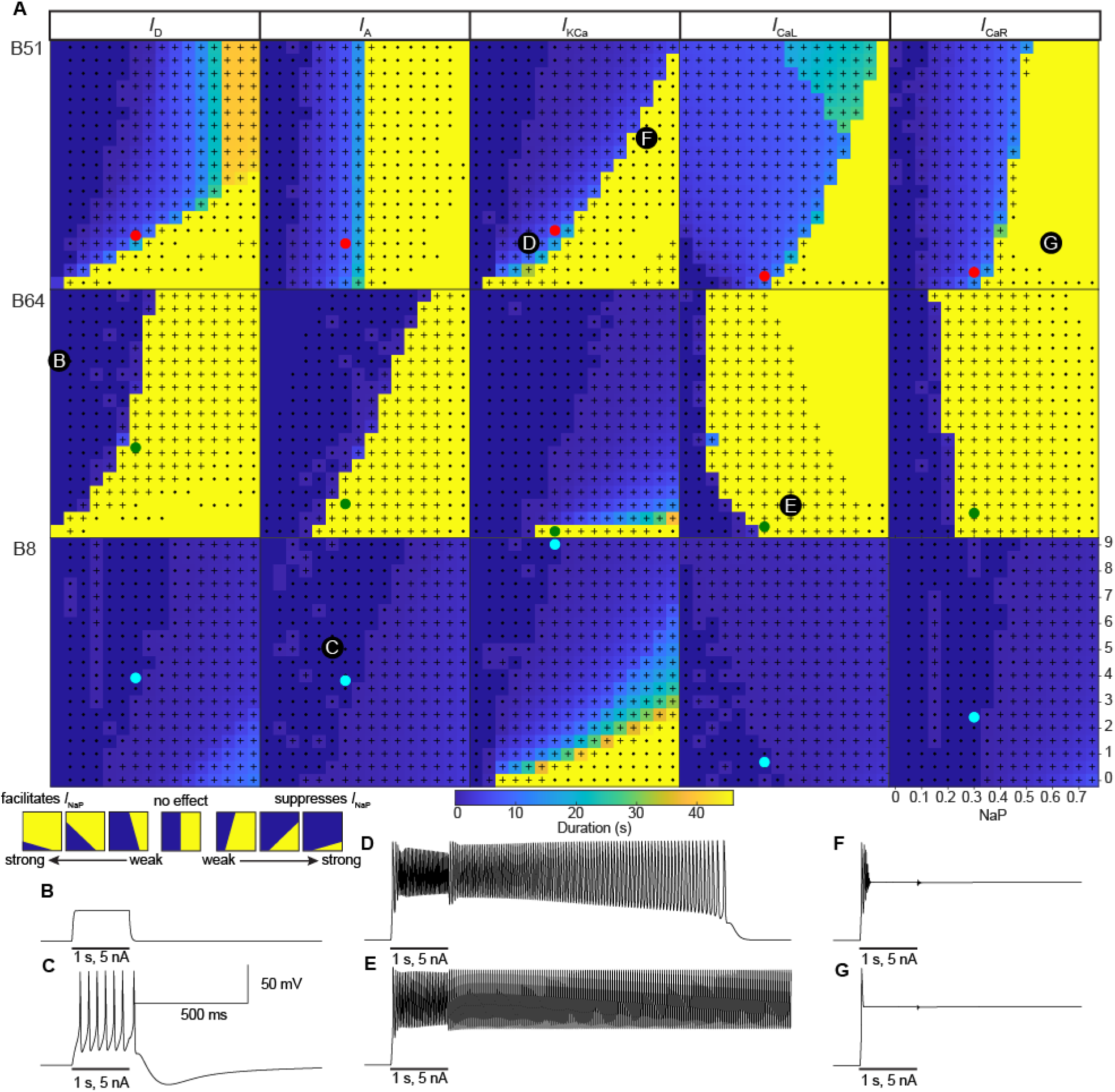
Heat maps of the parameter space indicate that ion channels affect each neuron in different ways. (A) Heat maps of the effects of varying g_max_ for *I*_D_, *I*_A_, *I*_KCa_, *I*_CaL_, and *I*_CaR_ vs. *I*_NaP_ on the plateau duration. The g_max_ of the outward currents and Ca^2+^ currents were each varied from 0 to 9 µS, while that of *I*_NaP_ was varied independently from 0 to 0.75 µS. The remaining model parameters were determined by experimental data presented in Figs. 1-5 and Supplemental Figs. 1-3. A guide to interpreting the maps is provided on the bottom left. The neuron model was held to -80 mV, similar to empirical experiments, and activity was elicited by a 1 s, 5 nA simulated positive current injection. The filled circles indicate the default values of g_max_ in the model (red = B51, green = B64, cyan = B8). The dots indicate a neuron that fires at least 3 action potentials. The + indicates action potentials occurred after the end of the positive current injection. (B – G) Illustrations of model responses. The g_max_ values for models in B – G are indicated in A. (B) Example of voltage response in a non-spiking model. (C) Example of a regular spiking model. (D) Example of a model with a self-terminating plateau potential. (E) Example of a spiking model with a plateau potential that does not self-terminate. (F) Example of a spike deficient model with a plateau that does not self-terminate. (G) Example of a model unable to elicit spikes with a plateau that does not self-terminate.

For the activation time constant:

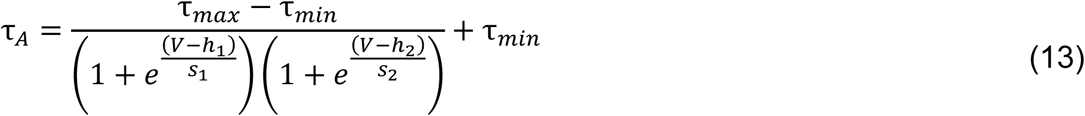

Where V is the voltage, h_1,2_ determine the voltage of the inflection points, s_1,2_ are slope parameters, τ _max_ is the maximum time constant, τ _min_ is the minimum time constant. For simplification and to aid in curve fitting of the current traces, the activation time constants of *I*_D_ and *I*_NaP_ were fixed to a single value, *I*_D_ was fixed to 0.01 s and *I*_NaP_ was fixed to 0.04 s. For voltage-gated Ca^2+^ channels, the activation time constants depended on the voltage clamp potential. To incorporate this voltage dependence into the model, the activation time constants were fit by hand with Eq. 13. The shapes of these curves in comparison to the time constant measurements are provided in Supplemental Fig. 2. For *I*_A_ (see Supplemental Fig. 3), the parameters for voltage dependency of activation were set to: B51, τ _max_ = 0.03 s, τ _min_ = 0.001 s, h = -30 mV, s = -5.7; B64, τ _max_ = 0.02 s, τ _min_ = 0.002 s, h = -20 mV, s = 5.7; B8, τ _max_ = 0.025 s, τ _min_ = 0.007 s, h_1_ = -20 mV, s_1_ = 6.

The inactivation, B_ij_ (Eq. 7), was defined by the following equations:

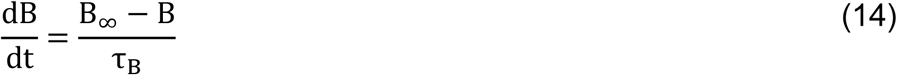

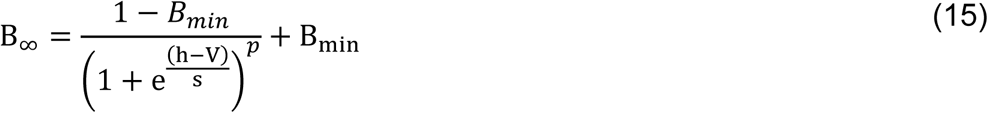

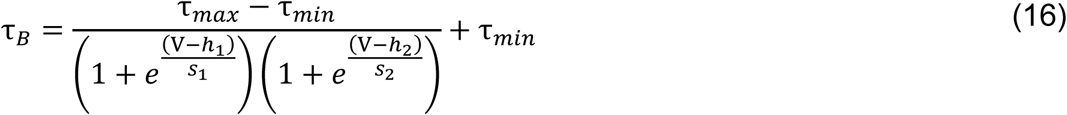

where V is the voltage, h_1,2_ determine the voltage of the inflection points, s_1,2_ are slope parameters, τ _max_ is the maximum time constant, τ _min_ is the minimum time constant. The level of inactivation (Supplemental Fig. 1) for each measured potential was fit with Eq. 15. For *I*_A_ and voltage-gated Ca^2+^ channels, *B*_*min*_ was fixed at 0. For voltage-gated Ca^2+^ channels, *h* was limited to negative voltages. The inactivation time constant was estimated by s in the exponential decay terms in Eq. 1–5. There didn’t appear to be any voltage dependence in inactivation time constants for *I*_A_ and *I*_CaL_ so the means of the inactivation time constants for all potentials were compared between the neurons (Supplementary Fig. 2A,D). There did appear to be voltage-dependence in the inactivation time constants for *I*_D_, *I*_CaR_, *I*_NaP_. For *I*_D_ and *I*_CaR_, the voltage dependence of the inactivation time constant seemed to be best fit by a sigmoidal equation and so the second term in the denominator of Eq. 16 was not included. The voltage dependence of inactivation time constants of *I*_NaP_ was fit using Eq. 16. The voltage dependence of inactivation time constants were fit by hand for *I*_D_, *I*_CaR_, and *I*_NaP_.

In SNNAP, a voltage-clamp experiment was simulated with the exact protocol as experimental procedures. The resulting simulated current response was then compared to empirical data obtained from the recordings (Supplementary Fig. 3). The parameters in Eqs. 12-13,15-16 and ḡvd_ij_ in Eq.7 were adjusted slightly to provide the best fit for the current trace and the voltage dependence of the time constants in Supplementary Fig. 2. This study did not characterize Na^+^-fast (*I*_Na_), so the model for this channel was guided by data of its characterization in *Aplysia californica* pedal neurons (Gilly et al., 1997) and was identical for all three neuron models.

### Statistical analyses

Matlab was used for all statistical analyses. For all comparisons *P* values < 0.05 were considered to be significantly different. A Lilliefors goodness-of-fit test was used to test for normality of the distribution. Non-parametric tests were used when normality could not be assumed. The Student’s t-test was used for comparing two groups and multiple groups were compared using ANOVA or Kruskal-Wallis test with the Tukey-Kramer post-hoc test.

## Acknowledgements

This work was supported by National Institutes of Health grant R01 NS101356. The authors thank P. Smolen for detailed comments on earlier drafts of the manuscript and E. Kartikaningrum for preparing cultures.

