## Supplemental Figures for "Genesis of bursting activity: Role of inhibitory constraints on persistent Na^+^ currents"

### SUPPLEMENTARY INFORMATION

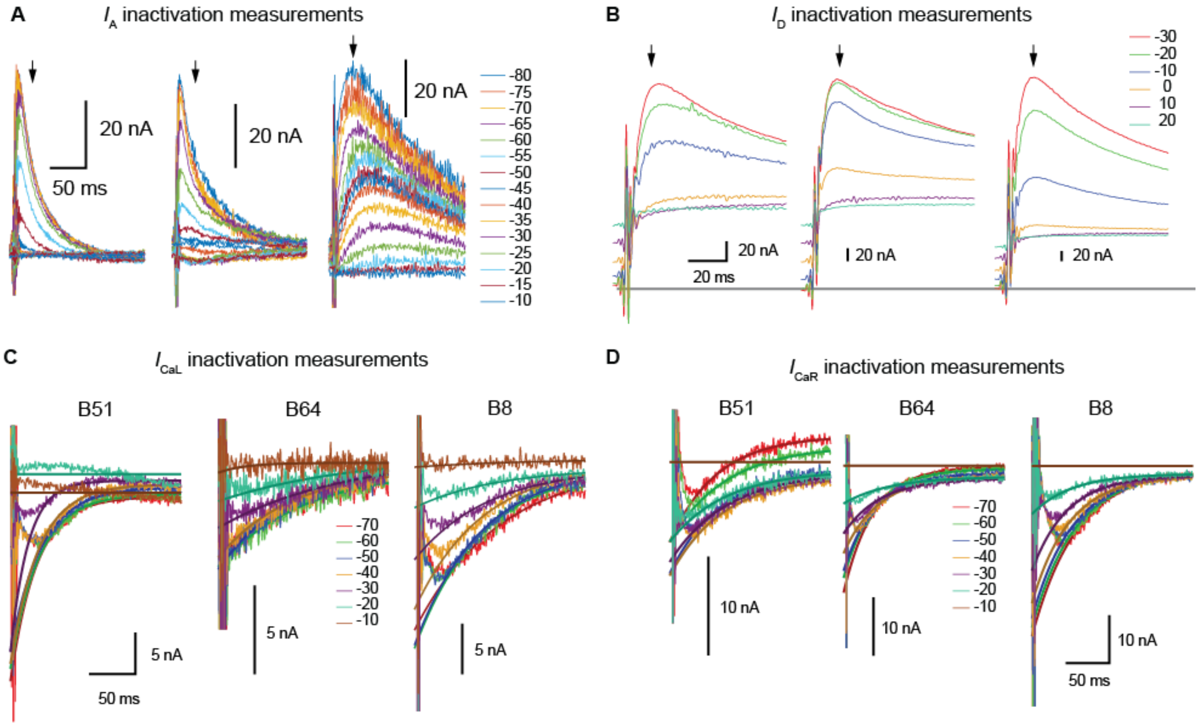


**Supplemental Figure 1. Protocol to measure inactivation.**

(A) Inactivation measurements for $I_A$. Currents measured at -10 mV elicited by a 1 s prepulse at the indicated potentials. Current was measured by taking the mean of a 4 ms segment 45 ms after start of pulse. Normalizing the measurements to the measured value with a -80 mV prepulse obtains the inactivation at each potential.

(B) Inactivation measurements for $I_D$. Currents measured at 30 mV elicited by a 1 s prepulse at the indicated potentials. Current was measured 18.7 ms after the start of the pulse. Normalizing the measurements to the measured value with a -30 mV prepulse obtains the inactivation at each potential.

(C) Inactivation measurements for $I_{CaL}$. Currents measured at 0 mV elicited by a 1.5 s prepulse at the indicated potentials. A single exponential was fitted to a 150 ms segment of data during the decay phase (starting 35 ms after the start of the pulse). The curve was then extrapolated back to 0 ms. Normalizing the value at 0 ms with that obtained from data with a -70 mV prepulse obtains the inactivation at each potential.

(D) Inactivation measurements for $I_{CaR}$. The same methods were used as in Panel C. For B51, a different set of cells were used than for Fig. 4K-S.

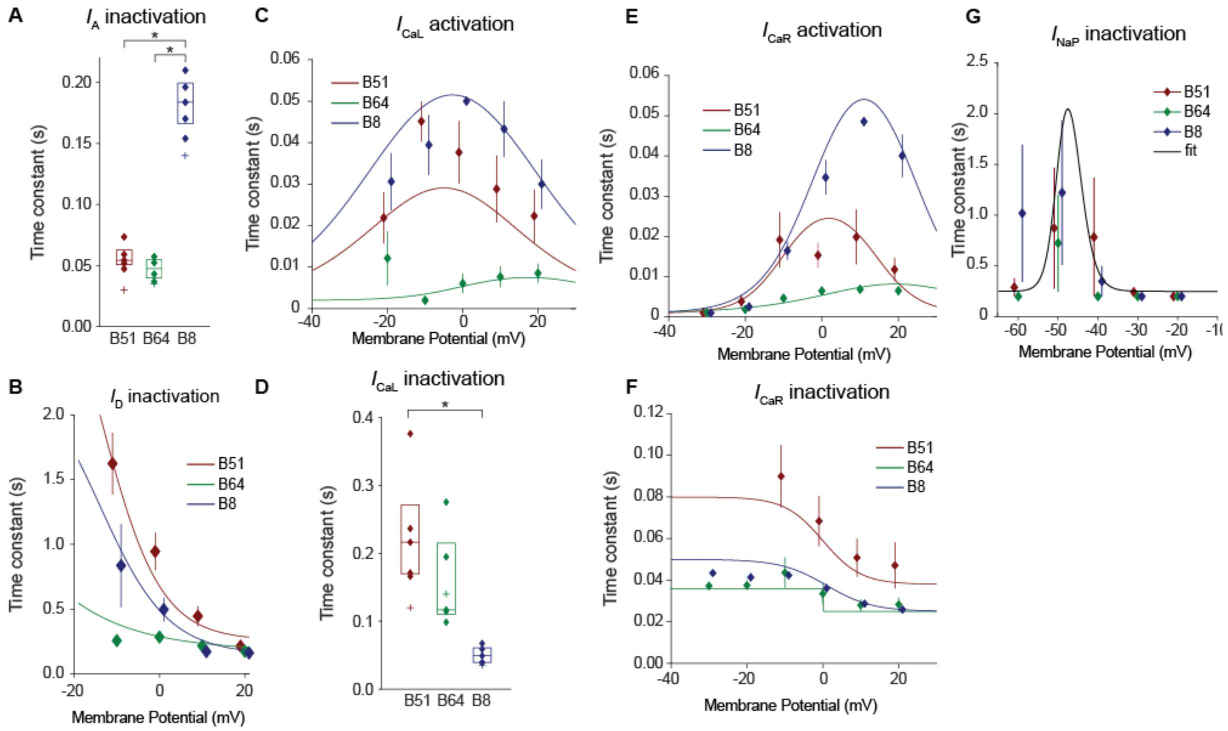


**Supplemental Figure 2. Comparison of empirically measured time constants and those used in the model.**

(A) The inactivation time constants for $I_A$ were averaged across all measured voltages. Inactivation time constant was greater in B8 $I_A$ compared to B51 and B64 (B51 = $0.054_{(0.051,\ 0.063)}$ MΩ, B64 = $0.048_{(0.040,\ 0.055)}$ MΩ, B8 = $0.18_{(0.17,0.20)}$ MΩ) ($F_{(2,13)}$ = 117.4, P = $3.8\text{x}10^{-8}$; *Post hoc* B51 vs. B64, P = 0.61; B51 vs. B8, P = $1.4\text{x}10^{-7}$; B64 vs. B8, P = $1.19\text{x}10^{-7}$). Plus sign indicates the values used for the model.

(B) The inactivation time constants for $I_D$. Clear voltage dependency can be seen. Eq. 16 without the second term in the denominator was used to fit the data. Solid lines indicate the time constants for the modeled current. Parameters were: B51, $\tau_{max}$ = 3 s, $\tau_{min}$ = 0.26 s, h = -14 mV, s = 7; B64, $\tau_{max}$ = 5.8 s, $\tau_{min}$ = 0.18 s, h = -60 mV, s = 15; B8, $\tau_{max}$ = 2 s, $\tau_{min}$ = 0.15 s, h = -14 mV, s = 8.

(C) The activation time constants for $I_{CaL}$. Clear voltage dependency can be seen. Eq. 16 was used to fit the data. Solid lines indicate the time constants for the modeled current. Parameters were: B51, $\tau_{max}$ = 0.06 s, $\tau_{min}$ = 0.002 s, $h_1$ = -15 mV, $s_1$ = -13, $h_2$ = 5, $s_2$ = 13; B64, $\tau_{max}$ = 0.0088 s, $\tau_{min}$ = 0.002 s, $h_1$ = 0 mV, $s_1$ = -8, $h_2$ = 35, $s_2$ = 8; B8, $\tau_{max}$ = 0.08 s, $\tau_{min}$ = 0.002 s, $h_1$ = -20.5 mV, $s_1$ = -13, $h_2$ = 15, $s_2$ = 13.

(D) The inactivation time constants for $I_{CaL}$ were averaged across all measured voltages. Inactivation time constant was greater in B51 $I_{CaL}$ compared to B8 and there was a trend towards being greater than B64 (B51 = $0.22_{(0.17,\ 0.27)}$ MΩ, B64 = $0.12_{(0.11,\ 0.22)}$ MΩ, B8 = $0.049_{(0.039,0.06)}$ MΩ) ($F_{(2,14)}$ = 9.75, P = 0.003; *Post hoc* B51 vs. B64, P =

0.22; B51 vs. B8, P = 0.0024; B64 vs. B8, P = 0.054). Plus sign indicates the values used for the model.

(E) The activation time constants for $I_{CaR}$. Clear voltage dependency can be seen. Eq. 16 was used to fit the data. Solid lines indicate the time constants for the modeled current. Parameters were: B51, $\tau_{max}$ = 0.03 s, $\tau_{min}$ = 0.001 s, $h_1$ = -10 mV, $s_1$ = -5.3, $h_2$ = 14, $s_2$ = 5.6; B64, $\tau_{max}$ = 0.02 s, $\tau_{min}$ = 0.001 s, $h_1$ = 13 mV, $s_1$ = -12.7, $h_2$ = 25, $s_2$ = 13; B8, $\tau_{max}$ = 0.076 s, $\tau_{min}$ = 0.001 s, $h_1$ = -1 mV, $s_1$ = -7.2, $h_2$ = 23, $s_2$ = 7.2.

(F) The inactivation time constants for $I_{CaR}$. Clear voltage dependency can be seen. Eq. 16 without the second term in the denominator was used to fit the data. Solid lines indicate the time constants for the modeled current. Parameters were: B51, $\tau_{max}$ = 0.08 s, $\tau_{min}$ = 0.038 s, h = 0 mV, s = 5; B64, $\tau_{max}$ = 0.036 s, $\tau_{min}$ = 0.025 s, h = 0 mV, s = 0.46; B8, $\tau_{max}$ = 0.043 s, $\tau_{min}$ = 0.025 s, h = 1 mV, s = 6.

(G) The inactivation time constants for $I_{NaP}$. Clear voltage dependency can be seen. Eq. 16 was used to fit the data. Solid lines indicate the time constants for the modeled current. Parameters for B51, B64, and B8 were: $\tau_{max}$ = 3 s, $\tau_{min}$ = 0.25 s, $h_1$ = -50 mV, $s_1$ = -2, $h_2$ = -45, $s_2$ = 2.

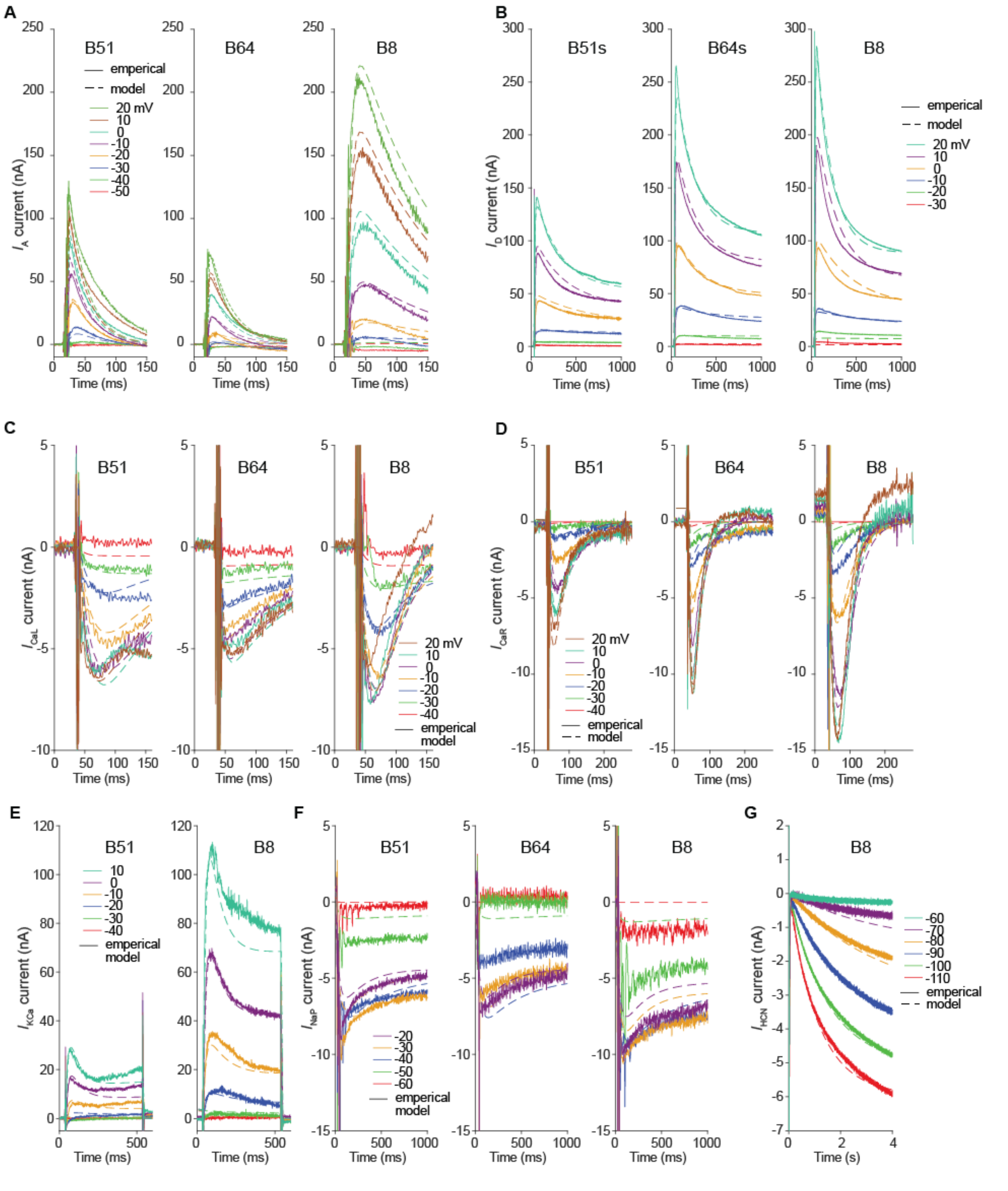


**Supplemental Figure 3. Comparison of voltage-clamp simulations and empirical data.**

(A) Mean current responses (solid line) from experiments measuring $I_A$ (Fig. 2A-I) for B51, B64 and B8 compared to simulations (dashed lines) of the same voltage-clamp protocol in the model neurons.

(B) Mean current responses of $I_D$ (Fig. 2K-T) for B51, B64 and B8 compared to simulation results.

(C) Mean current responses of $I_{CaL}$ (Fig. 4A-J) for B51, B64 and B8 compared to simulation results.

(D) Mean current responses of $I_{CaR}$ (Fig. 4K-S) for B51, B64 and B8 compared to simulations results.

(E) Mean current responses of $I_{KCa}$ (Fig. 3C-J) for B51, B64 and B8 compared to simulation results.

(F) Mean current responses of $I_{NaP}$ (Fig. 5C-J) for B51, B64 and B8 compared to simulation results.

(G) Mean current responses of $I_{HCN}$ for B51, B64 and B8 compared to simulation results.

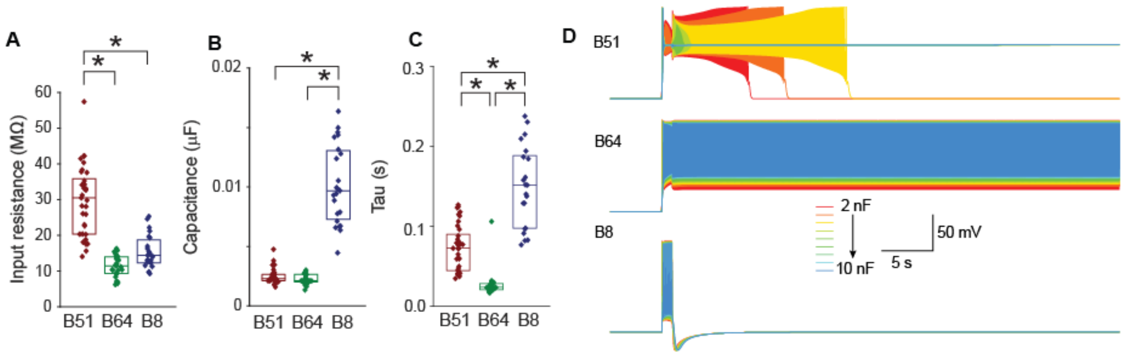


**Supplemental Figure 4. Differences in $R_{in}$ and capacitance cannot explain propensity to generate plateau potentials.**

(A) Input resistance measurements of B51, B64, and B8. Input resistance and time constant were measured by injecting a -1 nA current pulse. An exponential function was fitted to the first 1 s of the voltage trace during this current injection. Neuron B51 had a greater input resistance compared to B64 and B8 (B51 = $30.2_{(20.3,\ 35.8)}$ MΩ, B64 = $11.3_{(9.2,\ 13.8)}$ MΩ, B8 = $14.4_{(12.1,16.4)}$ MΩ) ($H_{(2)}$ = 50.7, P = $9.8 \times 10^{-12}$; *Post hoc* B51 vs. B64, P = $9.9 \times 10^{-10}$; B51 vs. B8, P = $1.4 \times 10^{-5}$; B64 vs. B8, P = 0.22).

(B) Capacitance measurements of B51, B64, and B8. Calculated by dividing the time constant by the input resistance. Neuron B8 had a greater capacitance compared to B51 and B64 (B51 = $0.0023_{(0.0021,\ 0.0027)}$ μF, B64 = $0.0022_{(0.0020,\ 0.0027)}$ μF, B8 = $0.0098_{(0.007,\ 0.013)}$ μF) ($H_{(2)}$ = 47.9, P = $4.0 \times 10^{-11}$; *Post hoc* B51 vs. B64, P = 0.51; B51 vs. B8, P = $1.3 \times 10^{-8}$; B64 vs. B8, P = $1.3 \times 10^{-9}$).

(C) Time constant measurements of B51, B64, and B8. B8 had a greater membrane time constant compared to B51 and B64 (B51 = $0.073_{(0.045,\ 0.090)}$ s, B64 = $0.024_{(0.021,\ 0.028)}$ s, B8 = $0.15_{(0.098,\ 0.19)}$ s) ($H_{(2)}$ = 62.1, P = $3.3 \times 10^{-14}$; *Post hoc* B51 vs. B64, P = $2.0 \times 10^{-6}$; B51 vs. B8, P = $6.0 \times 10^{-4}$; B64 vs. B8, P = $9.6 \times 10^{-10}$).

(D) The capacitance was varied from 2 nF (red, B51 empirical value) to 10 nF (blue, B8 empirical value). The models for neurons B51 or B64 were not converted to regularly firing neurons by increasing their capacitance. The models for neuron B8 was not converted to a model capable of generating a plateau by decreasing its capacitance.

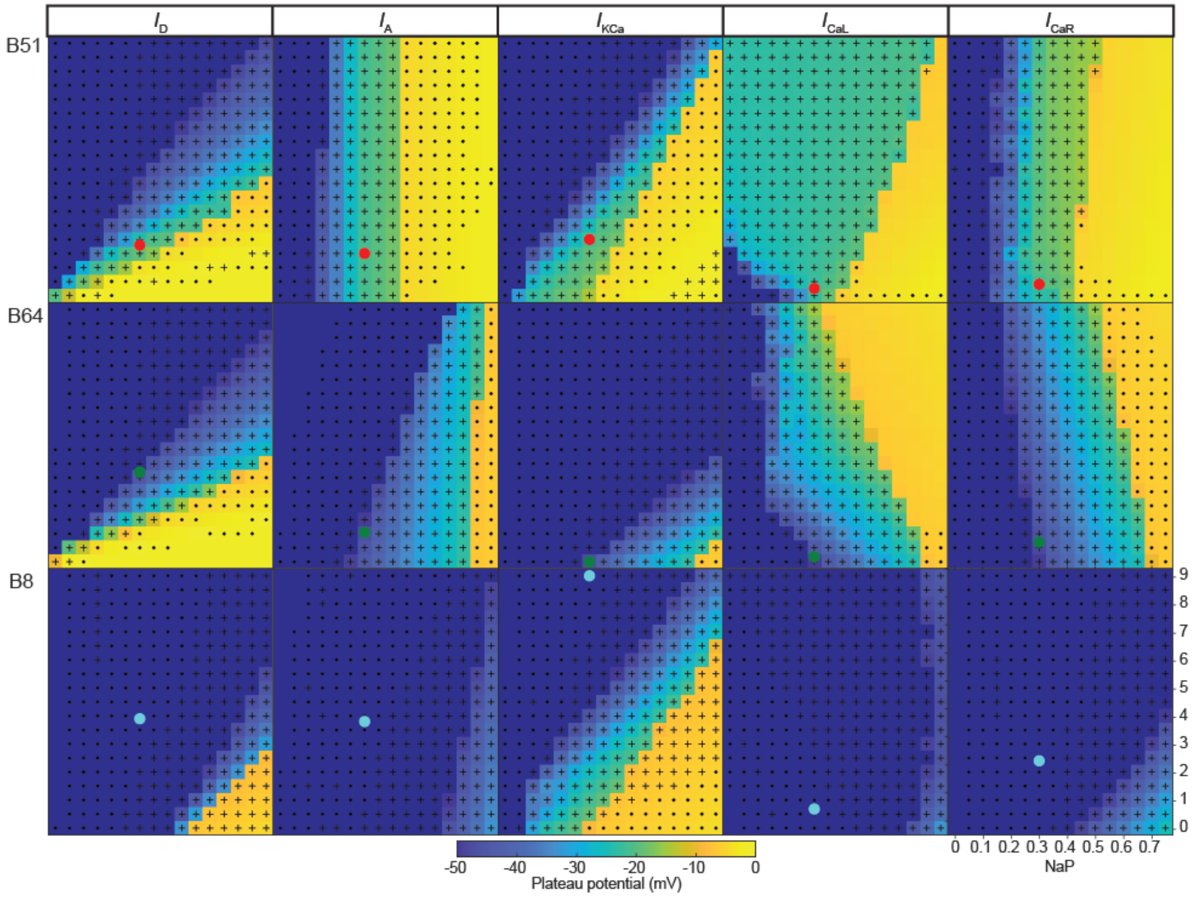


**Supplementary Figure 5. Heat maps of the parameter space indicate that ion channels differentially affect the depolarization of the plateau.**

(A) Heat maps of the effect of the $g_{max}$ parameter space for the $I_D$, $I_A$, $I_{KCa}$, $I_{CaL}$, and $I_{CaR}$ on the amount of depolarization of the plateau. The gmax of each channel was varied from 0 to 9 μS while $I_{NaP}$ was varied independently from 0 to 0.75 μS. The remaining model parameters were determined by experimental data presented in Figs. 1-5 and Supplemental Figs. 1-3. The neuron model was held to -80 mV similar to empirical experiments and activity was elicited by a 1 s, 5 nA simulated current injection. The dots indicate the default values of $g_{max}$ in the model (red = B51, green = B64, cyan = B8). Data from the same simulations as Fig. 7. Note the similarity with Fig. 7.

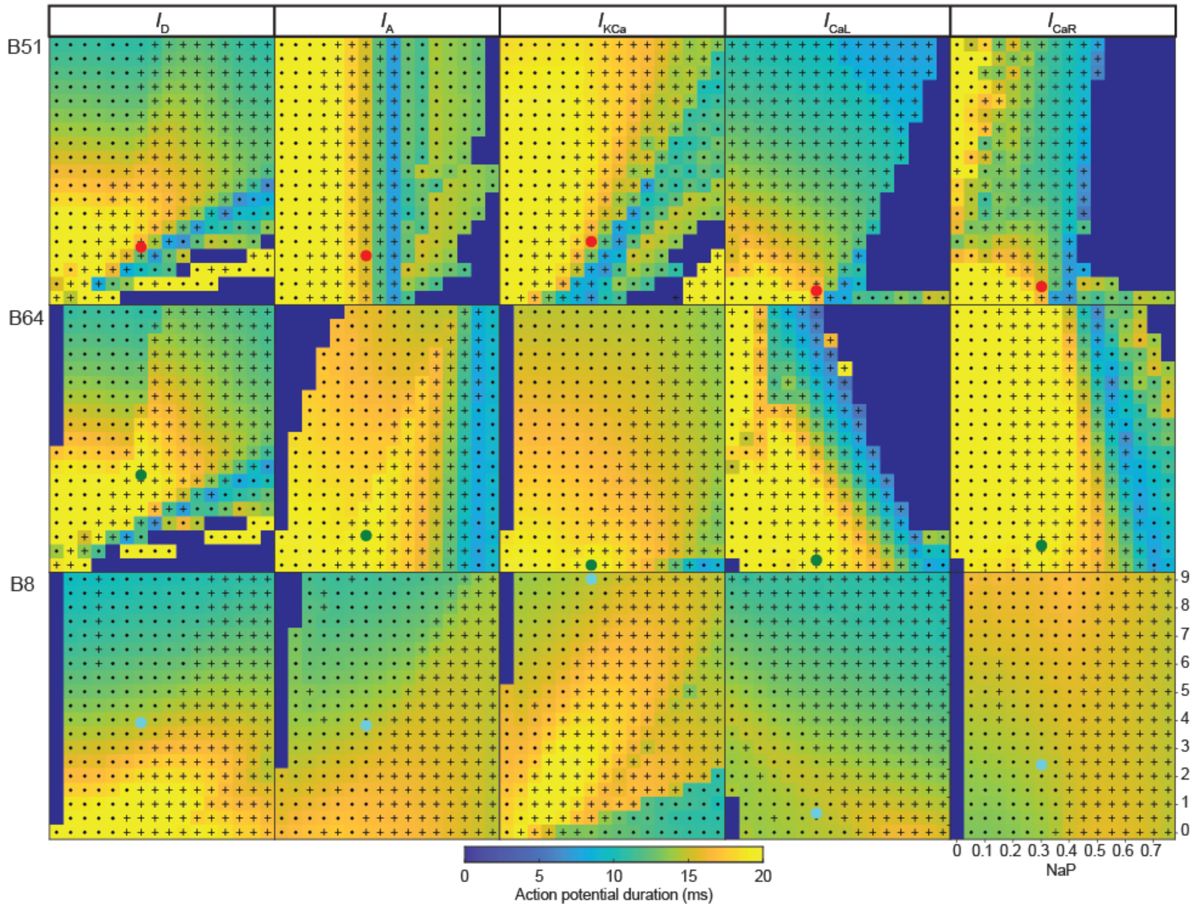


**Supplementary Figure 6. Heat maps of the parameter space displaying the complicated relationship between ion channels and the action potential duration.** Heat maps of the effect of the $g_{max}$ parameter space for the $I_D$, $I_A$, $I_{KCa}$, $I_{CaL}$, and $I_{CaR}$ on action potential duration. The gmax of each channel was varied from 0 to 9 μS while $I_{NaP}$ was varied independently from 0 to 0.75 μS. The remaining model parameters were determined by experimental data presented in Figs. 1-5 and Supplemental Figs. 1-3. The neuron model was held to -80 mV similar to empirical experiments and activity was elicited by a 1 s, 5 nA simulated current injection. The dots indicate the default values of $g_{max}$ in the model (red = B51, green = B64, cyan = B8). Data from the same simulations as Fig. 7.

| Channel | B51 | | B64 | | B8 | | | | B51 vs. B64 | B51 vs. B8 | B64 vs. B8 |
|---|---|---|---|---|---|---|---|---|---|---|---|
| | Model | Empirical | Model | Empirical | Model | Empirical | Statistics | Pvalue | | | |
| **$g_{max}$** | | | | | | | | | | | |
| $I_A$ | **1.5** | 1.02$_{(0.69, 1.1)}$ | **1.05** | 0.53$_{(0.51,0.54)}$ | **3.8** | 2.6$_{(1.8,3.1)}$ | $F_{(2,13)}$ = 23.7 | 0.0001 | 0.36 | $7.2x10^{-4}$ | $1.4x10^{-4}$ |
| $I_D$ | 1.8 | 1.6$_{(1.2, 2.0)}$ | 3.2 | 2.9$_{(2.3, 3.5)}$ | 3.9 | 3.5$_{(3.3, 4.1)}$ | $F_{(2,14)}$ = 9.19 | 0.0038 | 0.033 | 0.0034 | 0.43 |
| $I_{KCa}$ | 2 | 3.5$_{(1.9, 4.9)}$ | 0 | - | **9** | 13.6$_{(12.9, 27.5)}$ | $t_{(11)}$ = 4.52 | $8.7x10^{-4}$ | | | |
| $I_{CaL}$ | 0.25 | 0.25$_{(0.21, 0.28)}$ | 0.178 | 0.17$_{(0.17, 0.20)}$ | 0.68 | 0.68$_{(0.54, 0.94)}$ | $H_{(2)}$ = 10.26 | 0.0059 | 0.41 | 0.14 | 0.004 |
| $I_{CaR}$ | 0.4 | 0.34$_{(0.28, 0.38)}$ | **0.7** | 0.49$_{(0.40, 0.63)}$ | 2.4 | 2.03$_{(1.32, 2.5)}$ | $F_{(2,14)}$ = 31.0 | $1.12x10^{-5}$ | 0.74 | $1.61x10^{-5}$ | $7.61x10^{-5}$ |
| $I_{NaP}$ | 0.3 | 0.34$_{(0.25, 0.39)}$ | 0.3 | 0.23$_{(0.16, 0.42)}$ | 0.3 | 0.38$_{(0.29, 0.61)}$ | $F_{(2,24)}$ = 0.87 | 0.43 | | | |
| $I_{HCN}$ | | - | | - | 0.085 | | | | | | |
| **$h_{act}$** | | | | | | | | | | | |
| $I_A$ | -27 | -26.7$_{(-28.7, -25.0)}$ | -10.5 | -10.5$_{(-13.1, -6.7)}$ | -2.5 | -2.5$_{(-11.8, -0.72)}$ | $F_{(2,13)}$ = 22.5 | 0.0001 | 0.0014 | $1.4x10^{-4}$ | 0.50 |
| $I_D$ | **3** | -5.0$_{(-6.7, -3.0)}$ | 0.5 | -9.4$_{(-10.6, -3.1)}$ | 3 | -5.9$_{(-8.9, -2.8)}$ | $F_{(2,14)}$ = 0.31 | 0.74 | | | |
| $I_{KCa}$ | 0 | 4.1$_{(-8.9, 8.4)}$ | -1 | - | -1 | -1.7$_{(-4.8, 4.9)}$ | $t_{(12)}$ = -0.135 | 0.90 | | | |
| $I_{CaL}$ | -8 | -8.0$_{(-12.1, -5.9)}$ | -5 | -4.7$_{(-8.0, 0.05)}$ | **-14** | -3.7$_{(-7.6, -2.5)}$ | $F_{(2,14)}$ = 0.96 | 0.41 | | | |
| $I_{CaR}$ | 2 | 1.7$_{(1.4, 3.9)}$ | 2 | 3.9$_{(-3.5, 4.2)}$ | 2 | 2.0$_{(0.76, 3.9)}$ | $H_{(2)}$ = 0.04 | 0.98 | | | |
| $I_{NaP}$ | -45 | -47.1$_{(-49.9, -38.1)}$ | -45 | -39.5$_{(-44.3, -36.0)}$ | -45 | -44.7$_{(-50.8, -42.2)}$ | $F_{(2,24)}$ = 2.76 | 0.085 | | | |
| $I_{HCN}$ | | - | | - | -68 | | | | | | |
| **$s_{act}$** | | | | | | | | | | | |
| $I_A$ | 5.6 | 5.6$_{(4.6, 10.7)}$ | 11.24 | 11.2$_{(10.6, 13.3)}$ | 10 | 10.1$_{(9.7, 11.7)}$ | $F_{(2,13)}$ = 3.18 | 0.081 | | | |
| $I_D$ | **7.5** | 9.0$_{(8.3, 9.1)}$ | 7.7 | 9.2$_{(7.3, 9.9)}$ | **7.28** | 8.6$_{(7.4, 9.5)}$ | $F_{(2,14)}$ = 0.07 | 0.93 | | | |
| $I_{KCa}$ | 7.5 | 2.6$_{(0.55, 8.0)}$ | 8.5 | - | 8.5 | 7.9$_{(2.6, 10.7)}$ | $t_{(12)}$ = 0.576 | 0.58 | | | |
| $I_{CaL}$ | 7.89 | 7.9$_{(3.8, 8.8)}$ | 12.1 | 11.9$_{(10.1, 13.0)}$ | **6.5** | 7.9$_{(6.7, 13.0)}$ | $F_{(2,14)}$ = 1.71 | 0.22 | | | |
| $I_{CaR}$ | 7.25 | 7.2$_{(2.4, 8.5)}$ | 8.2 | 7.0$_{(6.7, 7.3)}$ | 5.7 | 4.8$_{(4.4, 5.5)}$ | $F_{(2,15)}$ = 1.67 | 0.226 | | | |
| $I_{NaP}$ | 2.35 | 2.2$_{(1.3, 3.3)}$ | 2.35 | 2.4$_{(1.9, 5.5)}$ | 2.35 | 2.7$_{(1.3, 5.5)}$ | $F_{(2,24)}$ = 0.39 | 0.68 | | | |
| $I_{HCN}$ | | - | | - | -4.8 | | | | | | |
| **$h_{ina}$** | | | | | | | | | | | |
| $I_A$ | -56 | -55.8$_{(-59.4, -55.8)}$ | -46.33 | -46.3$_{(-52.4, -44.5)}$ | -51 | -51.0$_{(-51.8, -45.9)}$ | $F_{(2,13)}$ = 5.94 | 0.018 | 0.029 | 0.036 | 0.95 |
| $I_D$ | -10.5 | -10.5$_{(-16.9, -9.3)}$ | -10.5 | -10.6$_{(-12.2, -2.7)}$ | -13 | -13.4$_{(-15.4, -9.4)}$ | $F_{(2,14)}$ = 1.57 | 0.25 | | | |
| $I_{KCa}$ | -25 | | -45 | - | -45 | | $t_{(12)}$ = 0.576 | 0.58 | | | |
| $I_{CaL}$ | -22 | -22.2$_{(-23.8, -19.2)}$ | -26.5 | -26.5$_{(-27.9, -25.5)}$ | -34.4 | -34.4$_{(-35.7, -30.0)}$ | $H_{(2)}$ = 11.1 | 0.0040 | 0.18 | 0.0026 | 0.27 |
| $I_{CaR}$ | -18.2 | -18.1$_{(-18.4, -18.1)}$ | -25 | -25.1$_{(-26.7, -22.9)}$ | -25 | -25.3$_{(-26.4, -23.3)}$ | $H_{(2)}$ = 9.38 | 0.0092 | 0.024 | 0.020 | 0.997 |
| $I_{NaP}^{\#}$ | -80 | - | -80 | - | -80 | - | | | | | |
| $I_{HCN}$ | - | - | - | - | - | - | | | | | |
| **$s_{ina}$*** | | | | | | | | | | | |
| $I_A$ | -4.3 | -4.3$_{(-5.0, -3.9)}$ | -7.38 | -7.4$_{(-8.1, -6.5)}$ | -9.4 | -9.4$_{(-12.2, -7.4)}$ | $F_{(2,13)}$ = 10.4 | 0.0029 | 0.082 | 0.002 | 0.19 |
| $I_D$ | -4.6 | -4.5$_{(-4.8, -4.4)}$ | -4.5 | -4.5$_{(-4.8, -4.4)}$ | -5.7 | -5.7$_{(-5.8, -5.4)}$ | $H_{(2)}$ = 5.42 | 0.067 | | | |
| $I_{KCa}$ | -7 | | -7 | - | -7 | | | | | | |
| $I_{CaL}$ | -0.95 | -0.99$_{(-4.1, -0.91)}$ | -7.47 | -7.5$_{(-7.6, -6.8)}$ | -6.26 | -6.3$_{(-6.6, -6.0)}$ | $H_{(2)}$ = 4.5 | 0.105 | | | |
| $I_{CaR}$ | -1.5 | -1.49$_{(-1.21, -2.18)}$ | -6.71 | -6.7$_{(-5.54, -7.9)}$ | -5.95 | -5.95$_{(-5.39, -7.09)}$ | $F_{(2,14)}$ = 25.5 | $4.77x10^{-5}$ | $8.5x10^{-5}$ | 0.0002 | 0.82 |
| $I_{NaP}^{\#}$ | -5 | - | -5 | - | -5 | - | | | | | |
| $I_{HCN}$ | - | - | - | - | - | - | | | | | |
| **$B_{min}$** | | | | | | | | | | | |
| $I_D$ | 0.364 | 0.36$_{(0.29, 0.42)}$ | 0.4 | 0.40$_{(0.37, 0.44)}$ | 0.274 | 0.27$_{(0.26, 0.31)}$ | $F_{(2,14)}$ = 5.43 | 0.021 | 0.33 | 0.21 | 0.016 |

**Supplementary Table 1. Table of all of the steady state parameters of the voltage gated ion channels.**

The parameter name is listed in the left column $g_{max}$ ($\bar{g}vd_{ij}$ in Eq. 7), $h_{act}$ (*h* in Eq. 12), $s_{act}$ (*s* in Eq. 12), $h_{ina}$ (*h* in Eq. 12), $b_{ina}$ (*s* in Eq. 12), $B_{min}$ ($B_{min}$ in Eq. 15). The parameter value used in the model is under the model column. The summary data of the measured value is in the empirical column. The summary data is represented in boxplots in Figs. 2-5. The empirical values are represented as median (25$^{th}$ percentile, 75$^{th}$ percentile). The statistical results comparing the empirical data are provided in the statistics and pvalue column and the post hoc analysis are provided in right three columns.
